# A Translational Model of MASLD-Associated HFpEF Defines Mitochondrial Dysfunction and Cardiac Plasticity During Disease Progression and Regression

**DOI:** 10.64898/2026.02.26.708088

**Authors:** Souradipta Ganguly, Betul Gunes, Yusu Gu, Jorge Suarez, Gautam Gupta, Kei Ishizuka, Rabi Murad, Tatiana Kisseleva, Wolfgang Dillmann, Kirk Peterson, Eric Adler, David A. Brenner, Debanjan Dhar

**Affiliations:** Sanford Burnham Prebys Medical Discovery Institute, La Jolla, CA, USA; Department of Medicine, University of California, San Diego, La Jolla, CA, USA; Department of Surgery, School of Medicine, University of California, San Diego, CA, USA

**Keywords:** HFpEF, Metabolic Dysfunction associated steatotic liver disease (MASLD), MASH, LV Dysfunction, Lifestyle Modification, Disease Regression, Liver-Heart Axis

## Abstract

Metabolic dysfunction-associated steatotic liver disease (MASLD) and its progressive form, metabolic dysfunction-associated steatohepatitis (MASH), are strongly linked to cardiac dysfunction, particularly, heart failure with preserved ejection fraction (HFpEF), yet the mechanisms underlying this association remain unclear because robust integrative preclinical models are lacking and the liver and heart are rarely studied as a coordinated system. Here we show that *Alms1^-/-^* (Foz/Foz) mice fed a Western diet develop MASH with advanced liver fibrosis accompanied by a HFpEF-like phenotype characterized by left ventricular hypertrophy, impaired cardiomyocyte contractility, reduced β-adrenergic reserve, elevated BNP, and increased mortality despite ejection fraction >50%. Liver fibrosis emerged as a parameter strongly associated with the cardiac dysfunction. Remarkably, dietary reversal improved hepatic architecture, normalized cardiac function, and improved survival, revealing marked plasticity of the liver-heart axis. Analyses of left ventricular (LV) remodeling revealed mitochondrial dysfunction, altered substrate utilization, and extracellular matrix remodeling in the LV, with consistent concordance to human HFpEF transcriptomic signatures. Ultrastructural studies confirmed mitochondrial injury and sarcomeric disorganization, linking metabolic failure to impaired cardiomyocyte performance. Together, these findings identify mitochondrial dysfunction as an important feature of MASLD-associated HFpEF-like cardiac dysfunction and establish the Foz/Foz model as a powerful platform for investigating pathophysiological pathways linking metabolic liver disease to cardiac dysfunction and for testing mechanism-based therapeutic strategies.

**STRUCTURED ABSTRACT:** *Background:* Metabolic dysfunction associated steatotic liver disease (MASLD) and its advanced form, MASH, are closely linked to cardiac dysfunction, particularly heart failure with preserved ejection fraction (HFpEF). However, the mechanisms underlying MASLD-associated HFpEF and its reversibility remain poorly understood, largely due to the lack of robust preclinical models. Here, we established a translational model of MASLD-associated cardiac dysfunction that recapitulates the key features of human HFpEF. We applied functional and transcriptomic analyses of the left ventricle (LV) to define the pathways associated with cardiac dysfunction and its reversibility.

*Methods:* *Alms1^-/-^* (Foz/Foz) mice and wild-type littermates were fed normal chow (NC) or Western diet (WD) for up to 36 weeks (wk). Reversibility was modeled by switching WD-fed Foz/Foz mice at 12wk back to NC for 12wk. Cardiac assessment included echocardiography, invasive hemodynamics with dobutamine stimulation, histopathology, electron microscopy and isolated cardiomyocyte contractility. LV transcriptomes were profiled by bulk RNA sequencing and analyzed by differential expression and pathway enrichment.

*Results:* Foz/Foz mice on WD for 24wk developed metabolic syndrome and MASH with advanced liver fibrosis. Cardiac phenotyping showed LV hypertrophy, impaired cardiomyocyte contractility, reduced β-adrenergic reserve, elevated plasma BNP, and increased mortality while the ejection fraction was preserved (>50%), consistent with HFpEF. The progression of cardiac dysfunction was closely associated with liver fibrosis that developed during MASH. Switching WD-fed Foz/Foz mice at 12wk to normal chow diet reversed hepatic fibrosis, restored LV function, and reduced mortality, demonstrating plasticity of the liver-heart axis. LV transcriptomic analysis revealed that cardiac impairment in these mice was associated with mitochondrial dysfunction, altered substrate utilization, extracellular matrix remodeling, and metabolic stress; pathways that are similarly dysregulated in human HFpEF. Cardiac electron microscopy revealed swollen mitochondria with disrupted cristae, which improved following dietary intervention.

*Conclusions:* Mitochondrial dysfunction and fibroinflammatory remodeling are prominent features of MASLD-associated cardiac dysfunction. Reversal of hepatic and cardiac phenotypes with dietary intervention, together with elucidation of underlying pathways, establish the Foz/Foz model as a useful translational platform for studying liver-heart axis in MASLD. 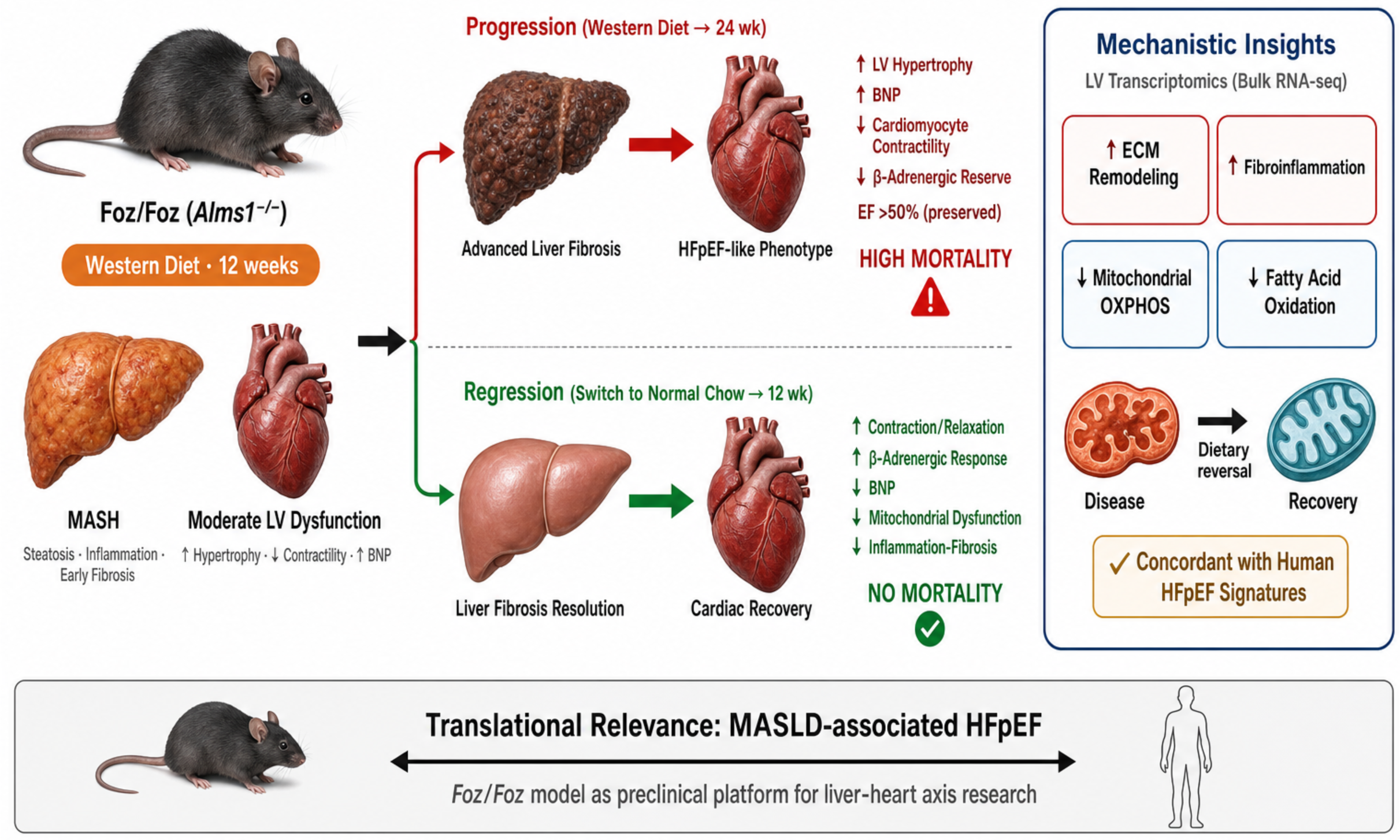

## 1. Introduction

Metabolic dysfunction-associated steatotic liver disease (MASLD) encompasses a spectrum from simple steatosis to metabolic dysfunction associated steatohepatitis (MASH), which can progress to fibrosis, cirrhosis, and hepatocellular carcinoma (HCC). MASLD affects more than 30% of adults worldwide, with prevalence rising each year[1–3]. As the hepatic manifestation of metabolic syndrome, MASLD is closely linked to obesity, insulin resistance, and dyslipidemia. Beyond the liver, MASLD exerts systemic effects, particularly on the cardiovascular system. Cardiovascular disease (CVD) is the leading cause of death in MASLD, followed by cancer and liver-related mortality[4, 5]. While both share metabolic and inflammatory pathways, the risk of CVD in MASLD is independent of traditional cardiovascular risk factors and increases with disease severity[6–8]. Recent work suggests two MASLD subtypes: a liver-specific form that progresses rapidly with limited CVD risk, and a cardiometabolic form marked by a higher incidence of CVD despite similar hepatic pathology[9]. Understanding why some patients develop cardiometabolic complications remains an important unmet need.

CVD in MASLD spans coronary artery disease, arrhythmias, stroke, and heart failure. Notably, up to 45% of MASLD patients exhibit left ventricular (LV) dysfunction, even when asymptomatic[10]. Although links between MASLD and atherosclerotic CVD are well-established, the mechanisms predisposing patients to LV dysfunction and heart failure remain unclear. Among cardiac outcomes, heart failure with preserved ejection fraction (HFpEF) is the most common and clinically significant in MASLD[11–13]. These patients show LV hypertrophy, altered geometry, and diastolic dysfunction, underscoring a strong liver-heart axis that is still poorly defined[14].

Progress has been hindered by the lack of animal models that recapitulate the integrated metabolic, hepatic, and cardiac features of human disease[15, 16]. In addition, the liver and heart are rarely examined together within the same study. The Foz/Foz mouse on a Western diet (WD) closely mirrors the multi-organ pathophysiology of metabolic syndrome[17], including obesity, insulin resistance, steatohepatitis, fibrosis, renal and cardiovascular dysfunction. Using this model, we show that cardiometabolic stress profoundly alters signaling, gene expression and remodeling of the heart. Notably, mitochondrial ultrastructure, metabolism, and bioenergetics were among the key pathways disrupted in LV during MASLD-associated HFpEF-like cardiac dysfunction.

Although lifestyle modification remains the cornerstone of MASLD management[18], most studies have focused primarily on hepatic outcomes, leaving the reversibility of MASLD-associated cardiac dysfunction largely unexplored. Here, we demonstrate that dietary reversal, switching MASH mice with advanced liver fibrosis and moderate HFpEF-like cardiac dysfunction from a Western diet (WD) to normal chow, restores hepatic health, attenuates cardiac dysfunction, and improves survival. Comprehensive transcriptomic mapping of LV gene signatures during both disease progression and resolution uncovered key molecular pathways associated with cardiometabolic remodeling across the MASLD spectrum.

## 2. Methods

### 2.1 Animal Models and Diet Protocols

Foz/Foz (*Alms1^−/−^*) mice on a C57BL/6J background were generously provided by Dr. Geoffrey C. Farrell (Australian National University Medical School) and further refined in our lab[17, 19, 20]. 6-8 week (wk) old male and female Foz/Foz mice, along with WT littermates, were fed either a Western diet (WD; AIN-76A, Test Diet, St. Louis, MO) containing 40% fat, 15% protein, 44% carbohydrates, and 0.2% cholesterol, or a standard chow diet (NC;12% fat, 23% protein, 65% carbohydrates) for up to 36wk. For dietary intervention studies, mice maintained on WD for 12wk were switched to chow for an additional 12wk (regression). Animals were randomly assigned to diet groups, with the individual mouse as the experimental unit. Mice were housed in pathogen-free conditions in individually ventilated cages with autoclaved food and water and maintained under a 12h/12h light/dark cycle. All procedures were conducted according to NIH guidelines and approved by the UCSD and SBP Institutional Animal Care and Use Committee (IACUC protocol #S07022 and #24-058 respectively).

### 2.2 Invasive Hemodynamic Analysis

A 1.4F micromanometer catheter (Millar Inc., Houston, TX) was inserted retrograde into the aorta via the left carotid artery and advanced into the LV in anesthetized mice (100mg/kg of ketamine and 10mg/kg of xylazine), intraperitoneal (IP) that are intubated (100-110 strokes/minute, 0.04-0.05 ml/stroke volume). A femoral vein was cannulated with a stretched PE50 tubing for drug administration. Baseline pressure measurements were obtained. Then, graded dobutamine doses of 0.75, 2, 4, 6, and 8 µg/kg/min were delivered using an infusion pump (PHD 2000, Harvard Apparatus, Holliston, MA) for 3 minutes at each dose. Data were recorded before and after bilateral vagotomy. Measurements of LV hemodynamic parameters were recorded and analyzed using LabChart (ADInstruments, Inc., Colorado Springs, CO).

### 2.3 Isolation of adult ventricular cardiomyocytes

Calcium-tolerant adult cardiomyocytes were isolated from ventricular tissue of mice by standard enzymatic digestion. Briefly, isolated hearts were perfused using a Ca^2+^-free Tyrode solution containing (in mM) 126 NaCl, 4.4 KCl, 1.2 MgCl_2_, 0.12 NaH_2_PO_4_, 4.0 NaHCO_3_, 10 HEPES, 30 2,3-butanedione monoxime, 5.5 glucose, 1.8 pyruvate, and 5.0 taurine (pH 7.3) for 5 min, followed by 0.9-1.0 mg/ml collagenase (type II, Worthington) for 20 min. Hearts were transferred to tubes containing fresh collagenase for an additional 10 min in a 37°C water bath. The heart tissue was mechanically dispersed and rinsed with gradually increasing extracellular calcium to 1mM. The cells were plated on 24×50-mm, no. 1 glass coverslips coated with laminin.

### 2.4 Measurement of Cardiomyocyte Contractility

Cardiomyocyte contractility was measured with a state-of-the-art integrated myocyte contractility workstation (IonOptix LLC). This method combines brightfield imaging with a high-speed camera recording system, along with sarcomere and cell-length detection algorithms, and application-specific analysis software to output reliable quantification of cardiomyocyte contractile function at the sarcomere level. Analysis outputs include determination of ±DL/Dt (dL/dt), representing peak shortening/re-lengthening velocities, and percent shortening for both sarcomere and cell length.

### 2.5 Statistical Analysis

Data are presented as mean ± SEM or as box-and-whisker plots (median, range, minimum/maximum), as indicated, and survival data are presented as Kaplan-Meier curves. For analyses involving transcriptomic deconvolution, group comparisons were performed using pairwise t-tests with Benjamini-Hochberg (BH) correction, and adjusted q-values are reported. For all other transcriptomic analyses, differential expression p-values were calculated using the Wald test and adjusted for multiple comparisons using the BH method. For all other datasets, group comparisons were performed using one-way or two-way ANOVA as appropriate, followed by Sidak’s multiple comparisons test when the overall ANOVA was significant. Survival analyses were performed using the log-rank (Mantel-Cox) test. Specific statistical tests used are indicated in the corresponding figure legends. P < 0.05 was considered statistically significant. Correlations were assessed using Spearman’s rank correlation. Sample sizes were not determined by formal a priori power calculations; instead, they were based on prior experience with this model and on effect sizes observed in our previous studies, where similar group sizes have consistently detected robust and biologically meaningful differences. These choices were further informed in consultation with a biostatistician to ensure appropriate study design and interpretation. All analyses were performed using GraphPad Prism (v10) and/or R (version 4.4.2; ‘stats’ package).

### 2.6 Data and code availability

All raw and processed sequencing data generated in this study have been deposited in the Gene Expression Omnibus (GEO) under accession number GSE304630, where they will be publicly accessible upon publication. Transcriptomic data from human HFpEF patients used for comparative analyses are publicly available in the European Nucleotide Archive (ENA) under accession number PRJEB62450. All analysis scripts and code used in this study will be released in a public repository at the time of publication to ensure full reproducibility. Additional methods are described in the supplementary materials.

## 3. Results

### 3.1 A pre-clinical model of Metabolic Syndrome (MetS), MASH and cardiometabolic dysfunction with mortality

*Alms1* mutated mice (Foz/Foz) serve as a preclinical model for studying the hepatic and extrahepatic manifestations of MetS (Fig. 1A-C)[17, 19, 21, 22]. Foz/Foz mice are hyperphagic (Fig. S1A) and when fed a high calorie, fructose and cholesterol-rich Western diet (WD), develop MASH and liver fibrosis by 12wk (Foz+WD 12wk)[17] accompanied by obesity, diabetes, hypercholesterolemia, and hypertriglyceridemia[17]. The disease progresses to advanced (stage 4) liver fibrosis (Fig. 1C) and about 75% mice develop HCC by 24wk[17]. Male Foz mice develop MASH with significant fibrosis, whereas female Foz mice, despite being hyperphagic and developing obesity, diabetes, dyslipidemia, liver injury, MetS, steatosis, and MASH[17] (Fig. 1B and Fig. S1B-F), remain resistant to liver fibrosis (Fig. 1C). Female mice are usually resistant to fibrosis development unless housed under thermoneutral conditions[23].

**Fig. 1:**
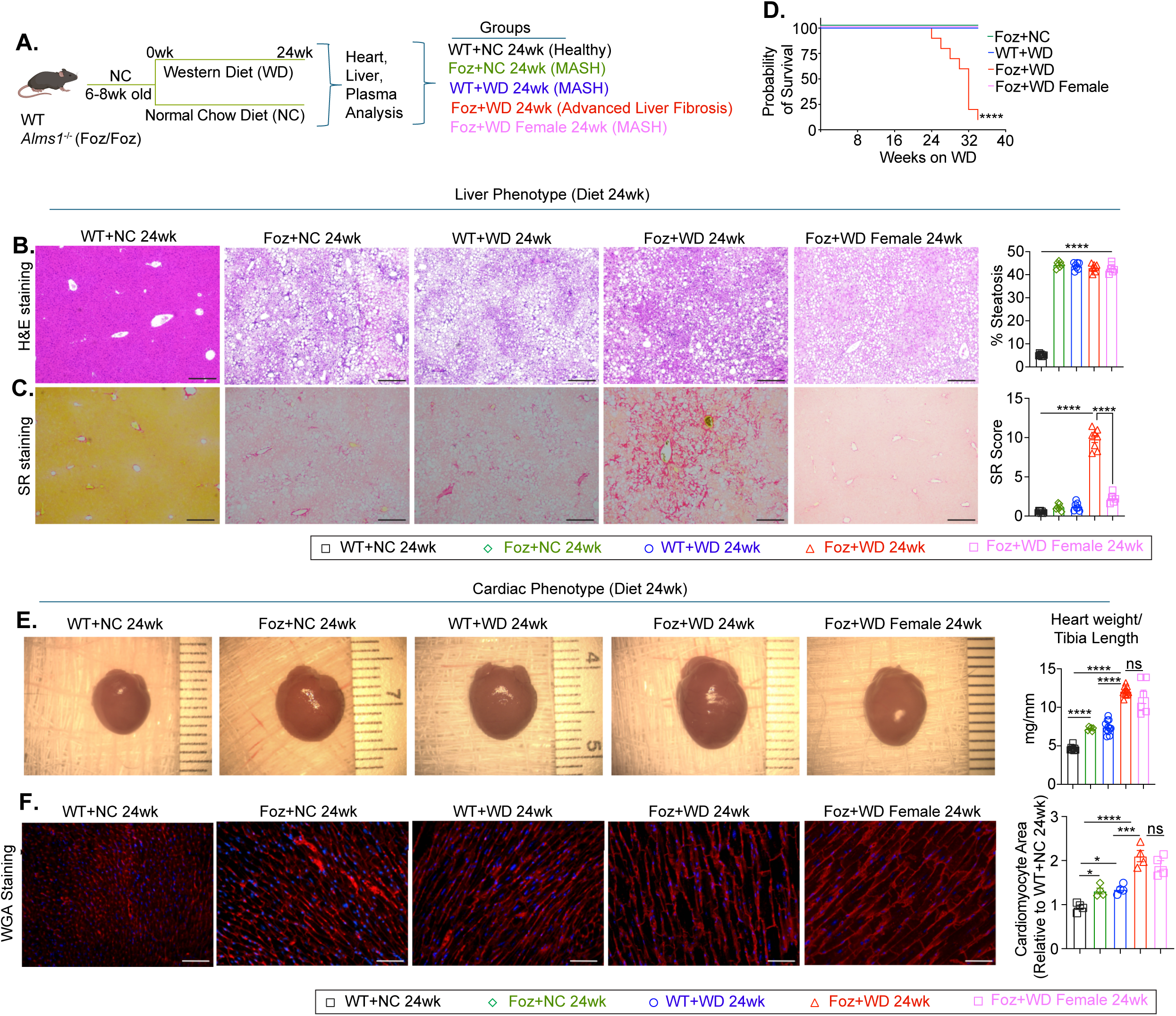
A pre-clinical model of MetS, MASH and cardiometabolic dysfunction with mortality. (A) Study design: 6-8wk old wild-type (WT) and *Alms1*^−/−^ (Foz/Foz) mice were fed either normal chow (NC) or Western diet (WD). At termination, heart, liver, and plasma samples were collected for analysis. (B-C) Representative hematoxylin and eosin (H&E) (B) and Sirius red (SR) (C) stained mouse liver sections with corresponding ImageJ quantifications. (scale bar: 260 µm). (D) Kaplan-Meier survival curve depicting the probability of survival over time (in weeks) for the indicated groups (sample size n=10 for each group). Survival curves were analyzed using the log-rank (Mantel-Cox) test. (E) Representative images of gross heart. Bar graph indicate heart weight normalized to tibia length. (F) Representative images of Wheat Germ Agglutinin (WGA) staining of cardiac sections (scale bar: 50 µm). Bar graph represents the ImageJ quantification of the cardiomyocyte area plotted relative to healthy controls (WT+NC 24wk). Bar plot data are presented as mean ± SEM. Sample size (n=4 mice per group) is indicated in each panel; each dot represents an individual mouse. Comparisons were performed using one-way ANOVA followed by Sidak’s multiple comparisons test. *p<0.05, **p<0.01, ***p<0.001, ****p<0.0001; ns, not significant.

WT mice on normal chow (WT+NC 24wk) showed no MetS, steatosis, or fibrosis, as expected (Fig. 1B, C and Fig. S1B-D). In contrast, age-matched Foz+NC mice developed hepatic steatosis with increased liver and body weight (Fig. S1B), hyperglycemia, dyslipidemia (Fig. S1C), and liver injury (Fig. S1D), indicating that prolonged hyperphagia alone can induce MetS and hepatic lipid accumulation. This phenotype resembled that of WT mice on a high calorie WD (WT+WD 24wk), and neither group developed fibrosis (Fig. 1B, C). These groups therefore represent MetS/MASH without liver fibrosis, in contrast to Foz+WD 24wk mice with MASH and advanced liver fibrosis.

Metabolic cage analysis revealed that Foz+WD 24wk mice showed reduced oxygen consumption, CO₂ production, and activity despite higher food and water intake, suggesting impaired energy metabolism (Fig. S1A). Intriguingly, mortality in Foz+WD male mice closely paralleled the progression of liver pathology, with deaths beginning around 24wk and reaching approximately 80% by 34wk (Fig. 1D), whereas all other groups exhibited 100% survival.

Despite increased liver damage in Foz+WD 24wk mice (Fig. S1D, S1G), liver function was preserved, as indicated by normal serum levels of bilirubin, albumin, and blood urea nitrogen (BUN), suggesting no decompensation to cause mortality (Fig. S1E). Consistently, necropsy findings did not reveal hepatic or renal failure as the primary cause of death (Fig. S1G). Given the clinical association between MASH, cardiac dysfunction (particularly HFpEF) and mortality in patients, we examined the hearts of these mice[11, 12, 15]. Compared to the healthy (WT+NC), all MetS groups (WT+WD, Foz+NC, and Foz+WD males and females) exhibited increased heart weight and cardiomyocyte size (Fig. 1E, F), with most severe hypertrophy in Foz+WD 24wk mice (Fig. 1E, F).

### 3.2 Left ventricular dysfunction in MASLD

To assess cardiac structural and functional changes associated with MetS and MASH and their correlation with liver fibrosis stage, we performed M-mode (Fig. 2A-J) and Doppler (Fig. S2A-C) echocardiographic analyses. M-mode echocardiography was used to measure cardiac structures, including chamber dimensions, heart size, and wall thickness, while Doppler echocardiography evaluated blood flow dynamics through the heart’s chambers and valves.

**Fig. 2:**
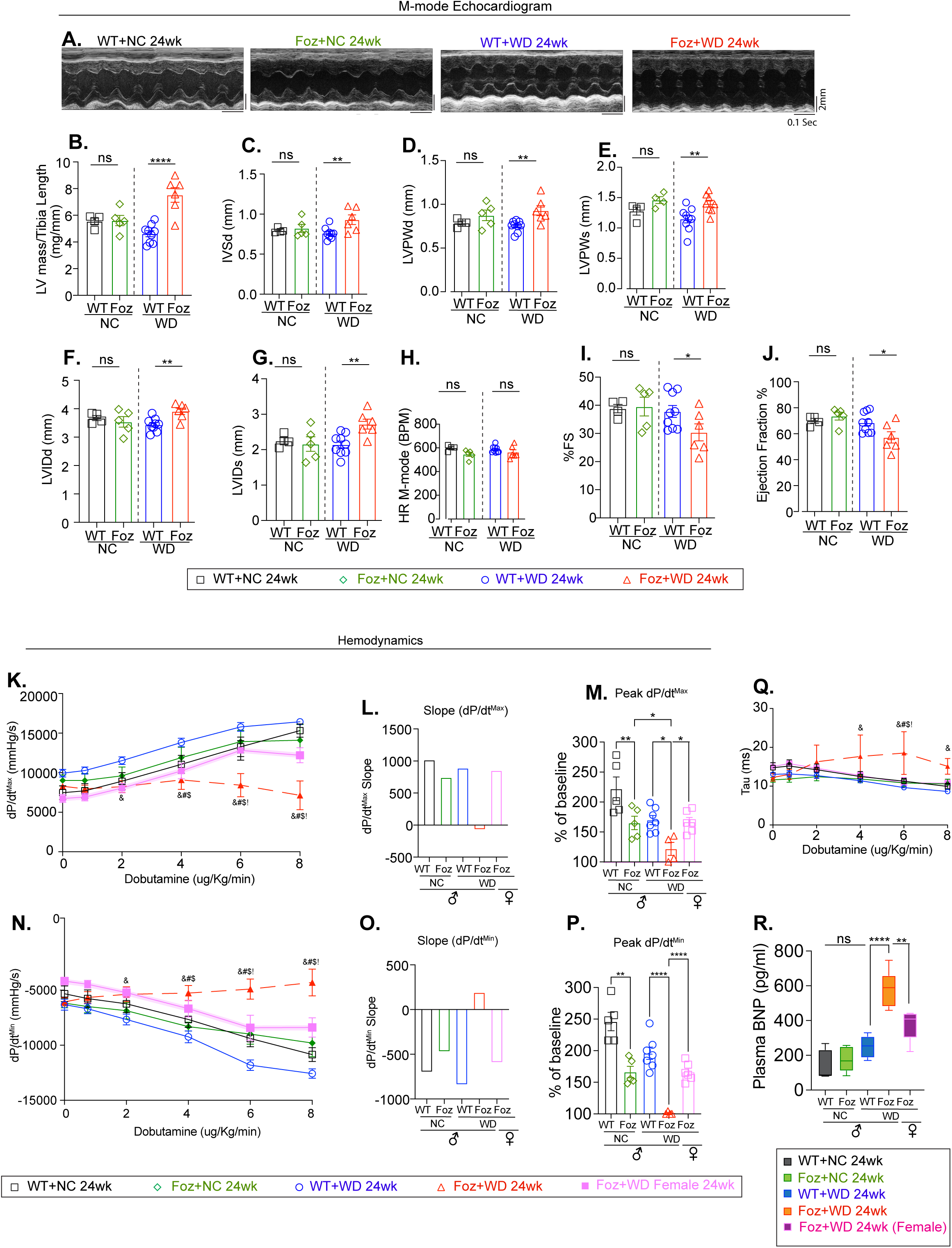
LV dysfunction in MASLD. (A) Representative M-mode echocardiographic images showing cardiac structural changes across groups. (B-J) Echocardiographic parameters were measured and plotted. (B) LV mass indexed to tibia length. (C) Interventricular septum thickness at diastole (IVSd) (D) LV posterior wall thickness at diastole (LVPWd), (E) LV posterior wall thickness at systole (LVPWs), (F) LV internal diameter at diastole (LVIDd), (G) LV internal diameter at systole (LVIDs) (H) heart rate (HR) (I) Fractional shortening (%FS), and (J) Ejection fraction (%EF), were measured. Sample size (n=4-9 mice per group) as indicated in each panel (each dot representing an individual mouse); analyzed by one-way ANOVA with Sidak’s multiple comparisons test when applicable. (K-Q) Hemodynamic assessment at baseline and with increasing dobutamine stimulation: (K-M) dP/dt^max^ (Maximal rate of LV pressure rise during contraction), shown as absolute values (K), slopes (L), and peak dP/dt^max^ expressed as a percentage of baseline (M). (N-P) dP/dt^min^ (Maximal rate of pressure decline during relaxation), with absolute values (N), slope (O) and peak dP/dt^min^ expressed as percentage of baseline (P). (Q) Relaxation constant Tau. Hemodynamic studies were analyzed either by one-way (for Fig. 2M, P) or two-way ANOVA (for Fig. 2K, N and Q) with Sidak’s multiple comparisons test. Sample size (n=4-7 mice per group); Symbols &, #, $, and ! denote P < 0.05 for the following comparisons: &: Foz+WD 24wk vs WT+WD 24wk; #: Foz+WD 24wk vs Foz+NC; 24wk $: Foz+WD 24wk vs WT+NC 24wk; !: Foz+WD 24wk male vs Female. (R) Plasma BNP levels were measured by ELISA and represented as a box-and-whisker plot, showing median, range, minimum/maximum values. All other data in this figure are presented as mean ± SEM. *P < 0.05, **P < 0.01, ***P < 0.001, ****P < 0.0001, ns, not significant.

LV mass increased by more than 30% in Foz+WD 24wk mice compared to other groups (Fig. 2B), accompanied by thickening of the interventricular septum (IVSd) and LV posterior wall (LVPWd and LVPWs), consistent with concentric hypertrophy observed in these mice (Fig. 2C-E). LV internal diameters were enlarged during both diastole (LVIDd) and systole (LVIDs) in mice with advanced liver fibrosis (Fig. 2F, G). Resting heart rate (HR) was similar across groups (Fig. 2H). Fractional shortening (FS), a key measure of systolic function, declined significantly exclusively in Foz+WD 24wk hearts (Fig. 2I), indicating impaired myocardial contractility. Ejection fraction (EF) was also reduced in Foz+WD 24wk (57%) mice versus other groups (69-74%) (Fig. 2J), yet remained above 50%, consistent with cardiac dysfunction exhibiting key features of HFpEF.

Interestingly, doppler echocardiography revealed no significant group differences in transmitral flow patterns or diastolic filling pressures (MV E/A and MV E/E’) (Fig. S2A-C), suggesting that LV dysfunction, is likely minimal or compensated at rest, a characteristic feature of HFpEF.

To further investigate the cardiac functional reserve under stress, we performed hemodynamic analysis under dobutamine (a β-adrenergic agonist) stimulation (Fig. 2K-Q Fig. S2D, E). The peak rate of contraction (dp/dt_max)_ and relaxation (dp/dt_min_) were measured. Dp/dt_max_ indicates the maximum rate of pressure increase during ventricular systole, reflecting myocardial contractile strength. Dp/dt_min_, denotes the maximum rate of pressure decline during diastole, measuring ventricular relaxation and compliance. These provide direct, load-sensitive insights into systolic and diastolic function[24–26].

In the healthy mice (WT+NC) dp/dt_max_ progressively increased with rising concentrations of dobutamine indicating preserved β-adrenergic responsiveness and functional cardiac reserve (Fig. 2K-M). Responses were modestly attenuated but remained largely preserved in the MetS+MASH groups without liver fibrosis (WT+WD and Foz+NC 24wk) (Fig. 2K-M). In stark contrast, Foz+WD 24wk mice showed no increase in dp/dt_max_ in response to dobutamine (Fig. 2K-M), demonstrating a complete loss of β-adrenergic responsiveness and severely impaired cardiac functional reserve, a hallmark of HFpEF. The peak dp/dt_max_ was highest in WT+NC, moderately reduced in the MetS+MASH mice without liver fibrosis and severely diminished in mice with advanced liver fibrosis (Foz+WD 24wk) (Fig. 2M). A similar trend was observed for dp/dt_min_, reflecting impaired diastolic function in mice with advanced liver fibrosis (Fig. 2N-P)[24–28]. This was further corroborated by a severely blunted lusitropic response, as the LV relaxation time constant, tau (ρ), failed to decrease during dobutamine infusion in the Foz+WD 24wk group (Fig. 2Q). Although heart rate increased in Foz+WD 24wk mice with rising dobutamine doses, maximal LV pressure failed to increase, consistent with the blunted dp/dt_max_ and dp/dt_min_ responses in these mice (Fig. S2D, E).

Interestingly, despite equivalent cardiac hypertrophy (Fig. S2F and Fig. 1F), Foz+WD 24wk females (MASH without liver fibrosis) had higher EF and FS (Fig. S2F) and superior dobutamine responses than males (Fig. 2K-P), paralleling improved survival (Fig. 1D). These findings suggest that liver fibrosis, rather than MetS or MASH alone, is more closely associated with LV impairment.

Plasma B-type natriuretic peptide (BNP) is a clinical marker of heart failure[29]. BNP levels were markedly elevated in Foz+WD 24wk males (∼600 pg/mL vs. <200 pg/mL baseline) (Fig. 2R), consistent with severe heart failure and increased mortality (Fig. 1D). WT+WD males and Foz+WD females exhibited intermediate BNP levels (Fig. 2R). Taken together, these data indicate that mice with advanced liver fibrosis (Foz+WD 24wk) develop combined systolic and diastolic dysfunction and heart failure. Although EF is reduced, it remains above 50% (Fig. 2J), consistent with features of HFpEF.

### 3.3 LV transcriptomics reveals key pathways associated with LV dysfunction

To define mechanisms underlying pathological cardiac remodeling, we performed bulk RNA sequencing (RNAseq) on isolated LVs from four groups (24wk WT+NC, Foz+NC, WT+WD, Foz+WD) (Fig. 3A). Unsupervised principal component analysis (PCA) identified three distinct transcriptomic clusters corresponding to their cardiometabolic phenotypes (Fig. 3B): (1) healthy (WT+NC), (2) MetS+MASH without liver fibrosis with mild hypertrophy (Foz+NC and WT+WD), and (3) MASH with advanced fibrosis and severe HFpEF-like LV dysfunction and (Foz+WD).

**Fig. 3:**
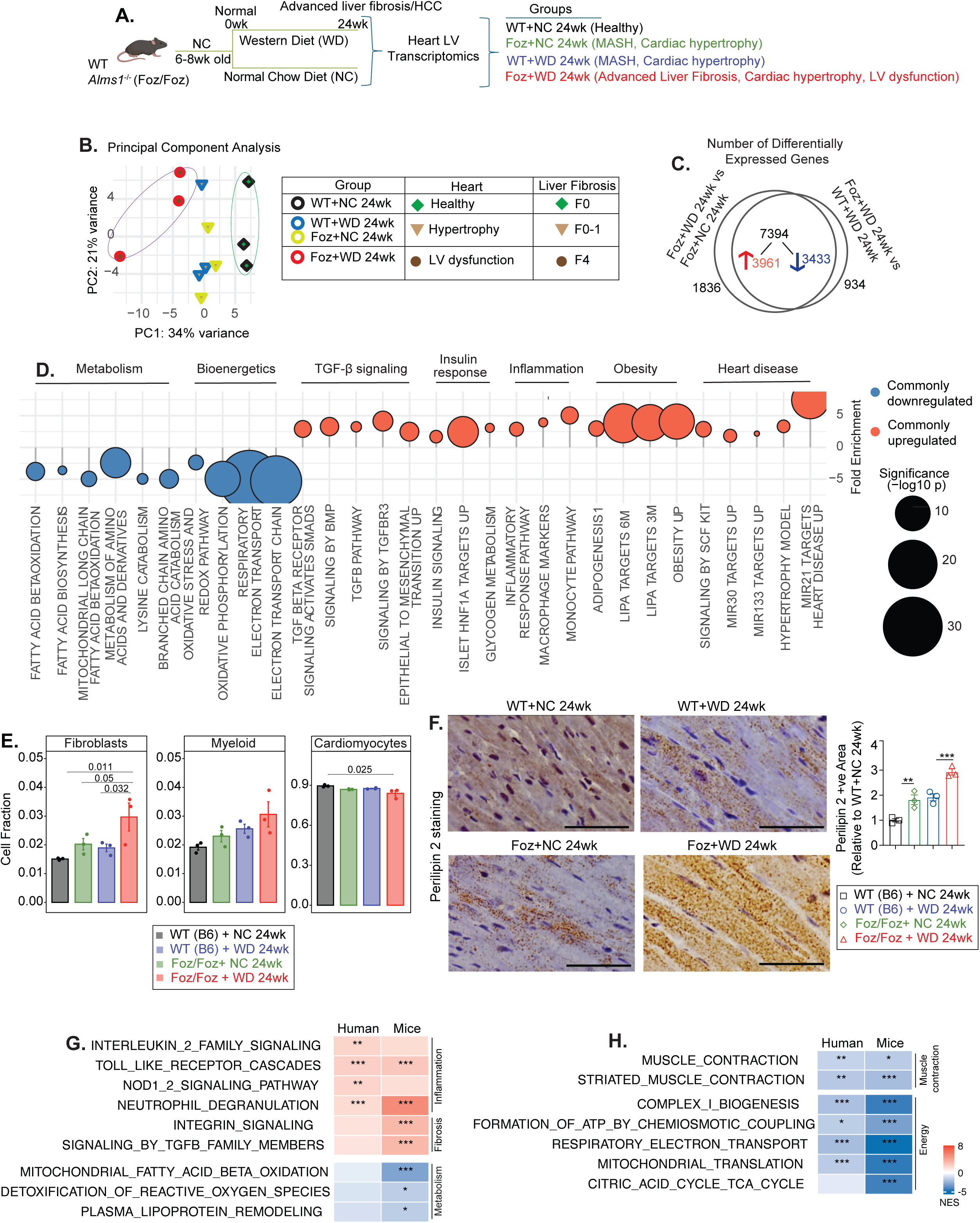
Transcriptomic analysis identifies molecular pathways associated with LV dysfunction. (A) Study design: WT and Foz/Foz mice were fed either NC or WD. 24wk post diet, LV was dissected, and total RNA was subjected to bulk RNAseq (n=3/group). (B) PCA plot illustrating the transcriptomic separation of samples and the corresponding liver and cardiac phenotypes. (C) Venn diagram showing 1,836 differentially expressed genes (DEGs) unique to Foz+WD vs Foz+NC, 934 unique to Foz+WD vs WT+WD, and 7,394 shared differentially expressed genes (3,961 upregulated; 3,433 downregulated) common to both comparisons. (D) Overrepresentation analysis of DEGs common to both comparisons, visualized as a bubble plot. Pathways overrepresented among upregulated (red) and downregulated (blue) genes are grouped by biological function; dot size reflects statistical significance (−log₁₀ P value), and the y-axis indicates fold enrichment. (E) Estimated cell-type proportions were inferred from bulk cardiac RNA-seq using CIBERSORTx-based deconvolution. Fibroblast, myeloid cell and cardiomyocyte fractions are shown across various groups. Group comparisons were performed using pairwise t-test with Benjamini-Hochberg (BH) correction, q values are indicated. (F) FFPE heart sections were subjected to IHC for perilipin 2. Representative images are shown (scale bar: 30µm). Bar plot shows imageJ quantification presented as mean±SEM with 3 mice/group. Group comparisons were performed by one-way ANOVA with Sidak’s multiple comparisons test. **p<0.01, ***p<0.001. (G-H) Gene set enrichment analysis (GSEA) comparing LV transcriptomic profiles from human HFpEF patients (ENA accession PRJEB62450) versus controls and Foz+WD 24wk versus WT+WD 24wk mice, with a heatmap displaying normalized enrichment scores (NES) for shared pathways enriched in both species.

Despite differences in genotype and diet, Foz+NC 24wk and WT+WD 24wk clustered together between the healthy and LV dysfunction groups. This suggests an intermediate cardiac phenotype marked by hypertrophy without progression to LV dysfunction. These two groups also shared metabolic and liver conditions limited to MASH (with little to no liver fibrosis; Fig. 1 and Fig. S1), reinforcing the liver-heart connection. Projecting the extent of liver fibrosis onto the cardiac PCA plot revealed a similar clustering pattern (Fig. 3B). These transcriptomic findings align with functional studies (Fig. 1, 2), demonstrating worsening cardiac dysfunction parallels MASH and liver fibrosis progression.

Differential expression analysis comparing Foz+WD with two non-fibrotic controls (Foz+NC and WT+WD) identified 1,836 diet-specific, 934 genotype-specific, and 7,394 shared differentially expressed genes (Fig. 3C) indicating convergence of dietary and genetic drivers. This approach allowed us to distinguish transcriptional changes linked specifically to pathological progression of HFpEF-like cardiac dysfunction, while separating the individual and combined influences of genotype and diet on the cardiac transcriptome. Downregulated genes (in Foz+WD 24wk) included key regulators of contractility (*Myh6, Tnnt2, Tnni3, Pln, Slc2a4*), fatty-acid β-oxidation (*Cpt1b, Acadm, Hadha*), amino-acid catabolism (*Bcat2, Mccc1, Mccc2*), and mitochondrial respiration (*Cox20, mt-Co1, Uqcrc2, Got2, Cpt2*) (Fig. S3A). In contrast, genes linked to insulin signaling, glycogen metabolism, inflammation, and fibrosis (e.g., *Col1a1, Tgfb1, Timp1, Vim*) were upregulated.

Overrepresentation analysis (ORA) of the 7,394 shared DEGs (Fig. 3D) revealed enrichment of TGF-β and insulin signaling, glycogen metabolism, inflammatory, obesity- and heart disease-related pathways (Fig. 3D, Fig. S3A) in Foz+WD 24wk group. Conversely, downregulated genes were primarily linked to fatty acid β-oxidation, amino acid catabolism, oxidative stress responses, and mitochondrial bioenergetic pathways, including oxidative phosphorylation and electron transport (Fig. 3D, Fig. S3A) in these mice. Gene set enrichment analysis (GSEA) comparing Foz+WD 24wk with Foz+NC and WT+WD groups corroborated these findings, showing negative enrichment of fatty acid metabolism and mitochondrial pathways and positive enrichment of fibrotic signaling (Fig. S3B-C) in Foz+WD 24wk group. Notably, while genes involved in fatty acid oxidation (FAO) and amino acid catabolism, the major energy generating processes in the healthy heart, were markedly repressed, genes associated with glycogen and glucose metabolism were upregulated (Fig. 3D, Fig. S3B-C) in Foz+WD 24wk heart. Together, these data indicate a metabolic shift toward altered substrate utilization, a hallmark of human HFpEF myocardium[30].

Cellular deconvolution of bulk RNAseq (Fig. 3E, Fig. S3E, F) revealed increased proportions of cardiac fibroblasts and myeloid cells in Foz+WD 24wk hearts, consistent with enhanced inflammatory and fibrotic signaling. Notably, despite this shift, the cardiomyocyte transcriptional footprint remained dominant (∼83-85%) across all conditions, in line with prior reports. Consistent with transcriptomic evidence of disrupted lipid metabolism pathways (Fig. 3D), Perilipin 2 immunostaining (a marker of intracellular lipid droplets) revealed pronounced lipid accumulation within cardiomyocytes of Foz+WD 24wk mice compared to controls (Fig. 3F). Notably, this lipid buildup occurred despite upregulation of LIPA (lysosomal acid lipase) pathways (Fig. 3D), suggesting a compensatory response to lipid overload or a failure in downstream lipid processing mechanisms. Sirius red staining revealed a modest increase in collagen deposition in Foz+WD 24wk heart compared with Foz+NC and WT+WD controls (Fig. S3D).

### 3.4 The Foz+WD Model Recapitulates Core Transcriptomic Features of Human HFpEF

To assess the transcriptomic relevance of this mouse model to human HFpEF, we compared pathway enrichment profiles in Foz+WD 24wk mice with those reported in LV from human HFpEF patients[31] (Fig. 3G, H). As metabolically relevant controls, we included WT+WD 24wk mice, which exhibit features of MetS and cardiac hypertrophy but retain intact β-adrenergic reserve.

Cross-species pathway analysis revealed concordance across inflammatory, metabolic, and contractile programs. Both human and mouse HFpEF hearts demonstrated upregulation of innate immune and inflammatory pathways, including Toll-like receptor, IL-2 family, and NOD1/2 signaling, as well as neutrophil degranulation, indicating conserved immune activation and supporting the translational fidelity of the Foz+WD model (Fig. 3G). Similarly, pathways governing oxidative phosphorylation, ATP production, the TCA cycle, and mitochondrial function were uniformly downregulated, reflecting shared deficits in myocardial energetics (Fig. 3H). Suppression of muscle contraction pathways in both species further suggests conserved impairments in sarcomere organization and mechanical performance. These cross-species similarities support the relevance of the Foz+WD model to human patients.

Taken together, these findings show that while MetS+MASH mice (without liver fibrosis) develop cardiac hypertrophy with preserved β-adrenergic responsiveness, progression to advanced liver fibrosis is accompanied by severe LV dysfunction, reduced survival, and transcriptomic signatures of fibrotic and inflammatory activation coupled with energetic insufficiency, marked by suppression of FAO and mitochondrial pathways, culminating in impaired myocardial contractility.

### 3.5 Effect of MASH-fibrosis regression on cardiac remodeling

Lifestyle management improves liver pathology and lowers cardiometabolic risk in human[32]. Previous research has demonstrated that dietary intervention significantly ameliorates MetS, MASH, liver fibrosis, and related signaling pathways in the liver[17, 19]. However, the effects of such interventions on cardiac remodeling and gene expression are unclear. To address this knowledge gap, two additional timepoints were incorporated into the study (Fig. 4A). (1) Early Cardiac Changes (Foz+WD 12wk): Foz+WD mice were examined at 12wk to capture moderate cardiac changes before severe LV dysfunction. At this stage, mice exhibit MASH with F2 liver fibrosis. (2) Dietary Intervention Cohort (Regression): A subset of Foz+WD 12wk mice was switched to a normal chow diet for another 12wk. This group was designed to model the effects of lifestyle modification and the potential for liver fibrosis resolution, mimicking clinical interventions in humans. Outcomes in this cohort were compared to both the Foz+WD 12wk (to evaluate pathways associated with disease reversal) and Foz+WD 24wk (to identify pathways associated with disease progression) groups.

**Fig. 4:**
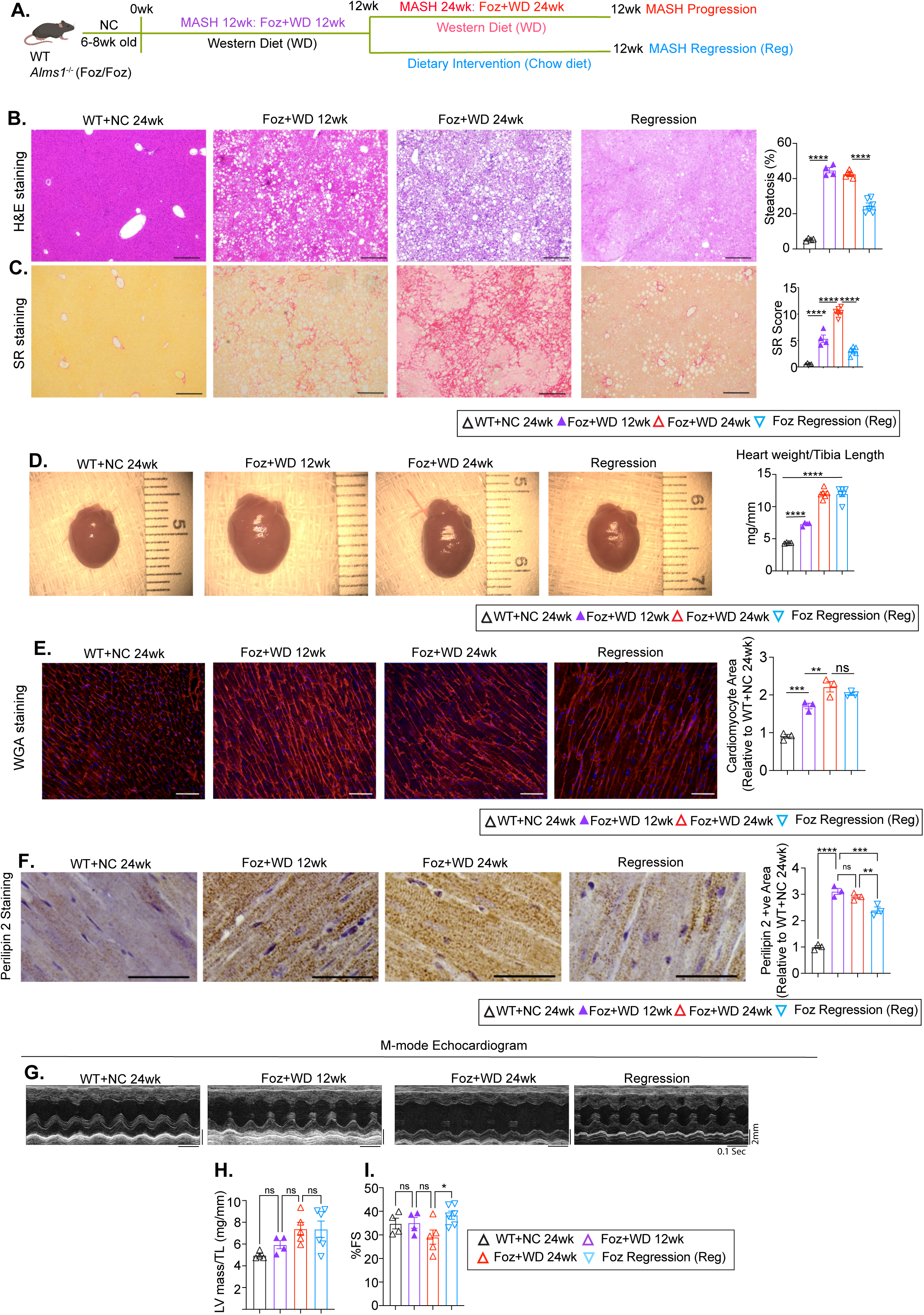
Effect of MASH-fibrosis regression on cardiac remodeling. (A) Study design. 6-8wk old Foz/Foz male mice were fed WD. Three Foz/Foz cohorts were established: (i) Foz+WD 12wk, (ii) Foz+WD 24wk, and (iii) Regression (Reg), where Foz mice were fed WD for 12wk and then switched to a normal chow diet for additional 12wk (Dietary intervention). WT+NC 24wk mice were used as healthy controls. (B, C) Liver pathology. FFPE liver sections were stained with H&E (B) and Sirius Red (SR) (C). (Scale bar: 260 µm). Bar plots represent imageJ quantifications. (D-I) Cardiac phenotype. (D) Gross images of hearts. Heart weight normalized to tibia length are plotted (right). (E) Representative images of WGA staining of cardiac sections. ImageJ quantification of cardiomyocyte area plotted relative to healthy control (WT+NC 24wk) is presented on the right. (F) FFPE heart sections were subjected to IHC staining for perilipin 2 (scale bar: 30 µm). Bar plots indicate ImageJ quantifications. (G-I) Before termination, mice were subjected to echocardiography. Representative M-mode echocardiography images from each group are shown illustrating cardiac structural changes. Selected echocardiogram parameters are plotted. (H) LV mass indexed to tibia length and (I) Fractional shortening (%FS). Bar plot data are presented as mean ± SEM. Sample sizes (n= 3-6 mice/group) are indicated in each panel (each dot representing an individual mouse). Group comparisons were performed using one-way ANOVA followed by Sidak’s multiple comparisons test, as applicable. *p<0.05, **p<0.01, ***p<0.001, ****p<0.0001, ns=not significant.

Foz/Foz mice develop steatosis within 1-2wk of WD, steatohepatitis by 4-6wk, and grade 2-3 fibrosis by 12wk[17]. Continued WD feeding to 24wk results in advanced liver fibrosis with progressive LV dysfunction and high mortality, as described above. Dietary intervention led to marked resolution of hepatic steatosis and fibrosis (Fig. 4B, C), along with other markers of liver damage and MetS (Fig. S4A-E). We examined whether this hepatic improvement was accompanied by transcriptomic and functional changes in the heart.

Heart weight and cardiomyocyte size increased progressively from healthy (WT+NC 24wk) to MASH (Foz+WD 12wk) and further to advanced liver fibrosis (Foz+WD 24wk) (Fig. 4D, E). During regression, cardiac hypertrophy persisted despite lower body and liver weight (Fig. 4D, E and Fig. S4A). Lipid accumulation in cardiomyocytes peaked at 12wk and remained high through 24wk (Fig. 4F). Dietary intervention led to a significant reduction in cardiac lipid content (Fig. 4F).

Echocardiography showed no difference in LV mass between progression and regression groups (Fig. 4G, H), indicating that dietary intervention did not reverse established cardiac hypertrophy. However, dietary intervention prevented progressive decline in fractional shortening in Foz+WD 24wk (compared to Foz Regression) (Fig. 4I). These findings suggest that while hypertrophy persisted, early dietary intervention preserved systolic function. No additional cardiac structural or functional changes were apparent between the progression and regression groups, by echocardiography (Fig. S4F-J).

### 3.6 Dietary intervention improves LV dysfunction and overall survival

Foz+WD 12wk mice (F2 liver fibrosis) exhibited attenuated β-adrenergic responsiveness compared to healthy controls (WT+NC) (Fig. 5A-F), whereas by 24wk (Foz+WD 24wk) this response was completely lost, as described above (Fig. 5A-F). Consistently, although Foz+WD 24wk mice showed impaired lusitropic reserve, with persistently elevated Tau during dobutamine stimulation, Foz+WD 12wk mice maintained a near-normal relaxation response comparable to WT+NC (Fig. 5G). These findings indicate that Foz+WD 12wk represents an early yet measurable stage of LV dysfunction (Fig. S5C).

**Fig. 5:**
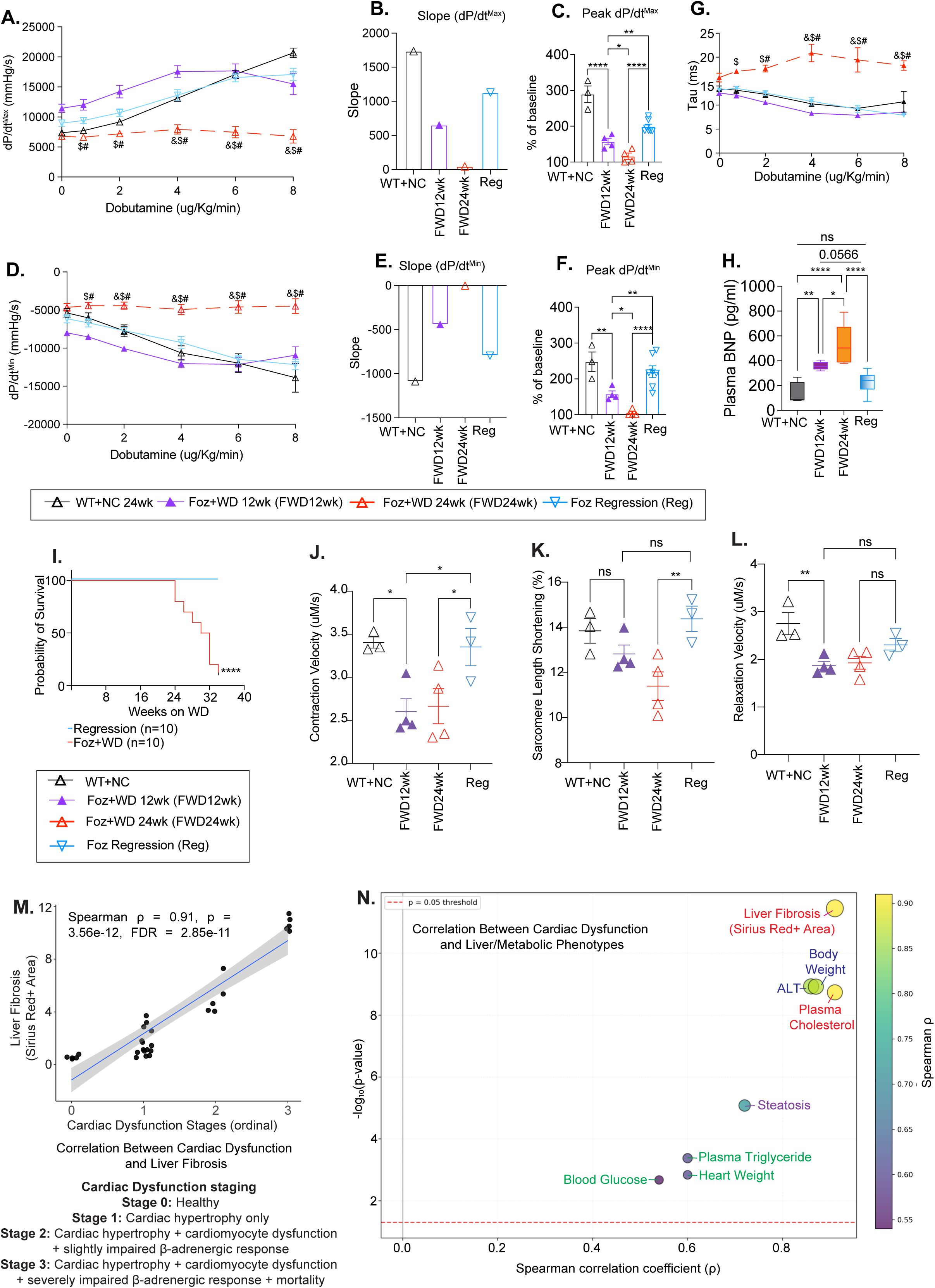
Dietary intervention improves LV dysfunction and overall survival. A subset of mice from groups described in Fig. 4A were subjected to the following analyses. (A-G) Hemodynamic indices of systolic and diastolic function measured under dobutamine stress. (A-C) dP/dt^max^ (Maximal rate of LV pressure rise during contraction), shown as (A) absolute values, (B) slopes, and (C) peak dP/dt^max^ expressed as a percentage of baseline. (D) dP/dt^min^ (Maximal rate of pressure decline during LV relaxation), with corresponding (E) slopes and (F) peak dP/dt^min^ plotted as percentage of baseline. (G) Relaxation constant Tau. Hemodynamic studies were analyzed by two-way ANOVA for Fig. 5A, D and G and one-way ANOVA for Fig 5C and F, with Sidak’s multiple comparisons test for both. Sample sizes (n) = 3-7 mice per group, Symbols, &, # and $ indicate P < 0.05 for the respective comparisons: &:Foz+WD 24wk vs WT+NC 24wk; #: Foz+WD 24wk vs Foz+WD 12wk; $: Foz+WD 24wk vs Regression 12wk. (H) Plasma levels of B-type natriuretic peptide (BNP) measured by ELISA and represented as a box-and-whisker plot, showing median, range, minimum/maximum values. (I) Kaplan-Meier survival curve depicting the probability of survival over time (in weeks) for the indicated groups (n=10). Survival differences were assessed using the Log-rank (Mantel-Cox) test. (J-L) Cardiac myocytes were isolated from hearts of the indicated groups and analyzed for the functional parameters including, (J) contraction velocity, (K) sarcomere length shortening, and (L) relaxation velocity. Sample size (n=3-4 mice per group) as indicated in each panel (each dot representing an individual mouse). These parameters were first measured in multiple (>100) isolated cardiomyocytes from each mouse and then averaged per mouse for statistical analysis. Group comparisons were performed using one-way ANOVA followed by Fisher’s LSD test. Data are presented as mean ± SEM unless otherwise mentioned. *p<0.05, **p<0.01, ***p<0.001, ****p<0.0001, ns: not significant. (M) Scatter plot showing the association between ordinal cardiac dysfunction stage and liver fibrosis burden quantified by Sirius Red positive area. Each point represents one animal. The solid line indicates linear regression with 95% confidence interval shading. (N) Summary bubble plot of Spearman correlations between cardiac dysfunction stage and liver/metabolic phenotypes. The x-axis shows Spearman correlation coefficient (ρ), and the y-axis shows −log₁₀ (p-value), bubble size represents −log₁₀(FDR) and the bubble color scale with correlation strength.

Importantly, dietary intervention initiated at 12wk not only halted further deterioration of cardiac function but also significantly improved β-adrenergic responsiveness, as reflected by steeper dp/dtmax and dp/dtmin slopes and higher peak values relative to both Foz+WD 12wk and 24wk groups (Fig. 5A-F). This intervention also preserved lusitropic function, with Tau values comparable to healthy controls (Fig. 5G), and was accompanied by improvements in heart rate and LV pressure (Fig. S5A, B).

Plasma BNP levels increased at 12wk (∼400pg/ml) and further at 24wk (∼500pg/ml) compared to the baseline (<200pg/ml) WT+NC group (Fig. 5H). Dietary intervention, however, not only prevented the increase in BNP levels, but also brought back the plasma BNP levels to near baseline (Fig. 5H). Importantly, dietary intervention conferred a profound survival benefit, achieving 100% survival compared to ∼20% survival (80% mortality) in age-matched WD-fed mice, underscoring the significant cardioprotective benefits of ameliorating MetS, MASH, and liver fibrosis (Fig. 5I).

We isolated cardiomyocytes and assessed contractility using sarcomere length measurements (SarcLem PMT software, IonOptix)[33]. Cardiomyocytes from both Foz+WD 12wk and 24wk mice demonstrated significantly reduced relaxation and contractile velocities, as well as decreased sarcomere shortening (Fig. 5J-L). Dietary intervention significantly improved contractile velocities (Fig. 5J), restored sarcomere shortening similar to the healthy controls (Fig. 5K), and relaxation velocities showed improvement trend compared to the Foz+WD groups (Fig. 5L) highlighting an adaptive enhancement of cardiomyocyte contractility during regression. Notably, despite similarly reduced contractile and relaxation velocities in isolated cardiomyocytes at 12 and 24wk (Fig. 5J-L), Foz+WD hearts at 12wk maintained a preserved LV response to dobutamine stress (Fig. 5A-G), suggesting preserved early functional reserve that is lost with disease progression.

Taken together, Foz+WD mice at 12wk represent an early stage of cardiac dysfunction, characterized by concentric LV hypertrophy, impaired cardiomyocyte contractility, reduced β-adrenergic responsiveness during dobutamine challenge, and moderately elevated plasma BNP levels. These abnormalities progress to a late-stage phenotype by 24wk, marked by complete loss of β-adrenergic reserve, markedly impaired active relaxation (prolonged Tau), and exacerbation of structural and functional deficits, ultimately culminating in increased mortality (Fig. 5I, S5C).

As we have characterized cardiac phenotypes across different mouse cohorts representing distinct stages of the MASLD continuum, we next performed a correlation analysis to assess the relationship between various metabolic parameters, hepatic and cardiac pathology (Fig. S5C). Quantitative analysis revealed a strong positive correlation between liver fibrosis severity and cardiac dysfunction (Spearman’s ρ=0.91, p=3.56 × 10⁻¹²), indicating tightly coupled progression of hepatic injury and cardiac impairment in this model (Fig. 5M, N and Fig. S5C).

### 3.7 LV remodeling pathways and gene expression changes during MASH progression

Understanding the signaling networks that govern disease progression and resolution is critical. To delineate the transcriptomic changes underlying the transition from healthy myocardium to early contractile impairment to severe LV dysfunction, we conducted gene expression analyses across three experimental cohorts: WT+NC 24wk (healthy), Foz+WD 12wk (F2 liver fibrosis) and Foz+WD 24wk (advanced stage 4 liver fibrosis) (Fig. 6A). PCA of the LV transcriptomes revealed a gene expression trajectory (red vector) indicative of progressive transcriptional reprogramming aligned with the exacerbation of cardiac dysfunction, that paralleled hepatic disease progression from healthy (WT+NC 24wk) to fibrotic MASH (Foz+WD 12wk) and ultimately advanced liver fibrosis (Foz+WD 24wk) (Fig. 6B).

**Fig. 6:**
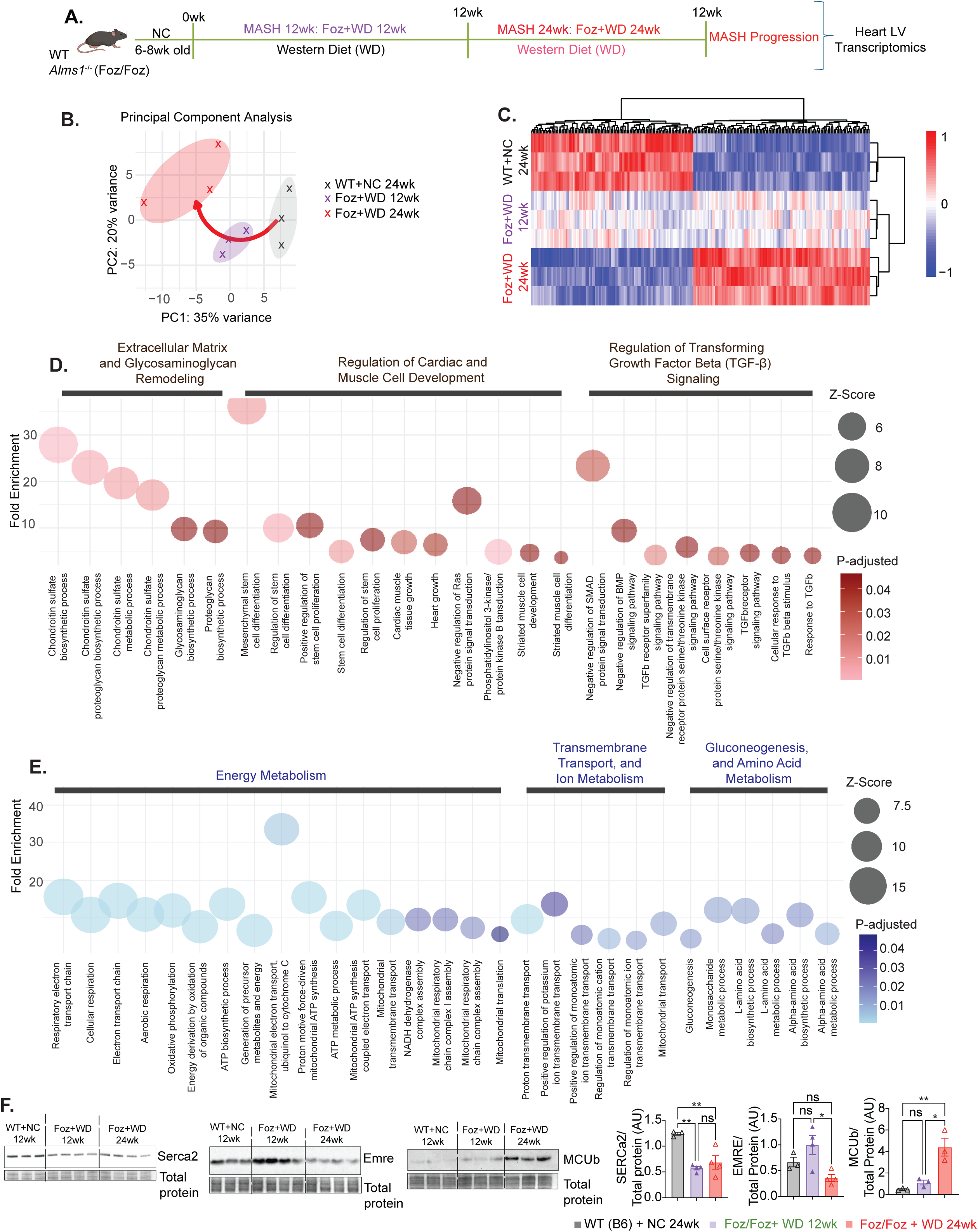
Molecular pathways underlying LV remodeling during MASH progression. (A) Study design: Foz/Foz mice were fed a Western diet (WD) for 12wk to induce MASH with moderate fibrosis. A separate cohort was maintained on WD for 24wk to promote progression to advanced fibrosis. (B) Principal Component Analysis (PCA) plot showing transcriptomic differences among the experimental groups. The red arrow indicates the progression from healthy WT+NC to intermediate dysfunction in Foz+WD 12wk, culminating in severe LV dysfunction in Foz+WD 24wk. (C) Heatmap of differentially expressed genes (DEGs) showing consistent expression changes across the spectrum of disease severity-from WT+NC 24wk (healthy), Foz+WD 12wk (intermediate), to Foz+WD 24wk (advanced dysfunction)-highlighting coordinated upregulation (red) and downregulation (blue) patterns. (D-E) Overrepresentation analysis of these coordinated upregulated (D) and downregulated (E) DEGs visualized as a bubble plot. Dot size is proportional to significance (-log10 p-value), and the y-axis displays fold enrichment. Pathways are grouped based on their biological relevance. (F) Representative immunoblots of SERCA2, EMRE, and MCUb in isolated cardiomyocytes from WT+NC (12wk) and Foz+WD (12wk and 24wk). Total protein staining is shown as a loading control. Densitometric quantification of SERCA2, EMRE, and MCUb normalized to total protein is presented on the right. Group comparisons were performed using one-way ANOVA followed by Sidak’s multiple comparisons test. *p<0.05, **p<0.01, ns=not significant.

To define molecular pathways driving the transition of mild to severe LV dysfunction, we performed supervised whole-transcriptome analysis focusing on genes with consistent directional changes across the three disease stages (Fig. 6C). The resulting heatmap revealed a clear transcriptional trajectory, with the LV transcriptome of Foz+WD 12wk (MASH-fibrosis) mice exhibiting an intermediate pattern between healthy and advanced liver fibrosis groups, suggesting a stepwise continuum of cardiac remodeling paralleling liver disease progression (Fig. 6C). Overrepresentation analysis of upregulated genes showed enrichment in extracellular matrix (ECM) remodeling, glycosaminoglycan processing, and TGF-β signaling, hallmarks of fibrotic activation and maladaptive structural remodeling (Fig. 6D) in Foz+WD 24wk group. Additional enrichment in “cardiac muscle tissue growth” and “regulation of stem cell proliferation” pathways may reflect compensatory adaptations to chronic stress in these mice. In contrast, downregulated genes were enriched for mitochondrial metabolism and energy production pathways, consistent with impaired mitochondrial efficiency and bioenergetic decline (Fig. 6E) in Foz+WD 24wk mice. Suppression of pathways such as “positive regulation of potassium ion transmembrane transport” and “mitochondrial ATP synthesis” further indicates disrupted ionic homeostasis and mitochondrial dysfunction, hallmarks of failing myocardium in these mice.

GSEA of pairwise comparisons of the above groups revealed that, in Foz+WD 12wk hearts, pathways related to fatty acid oxidation and mitochondrial bioenergetics (e.g., ETC and oxidative phosphorylation) were negatively enriched relative to healthy mice (WT+NC 24wk) but positively enriched compared to Foz+WD 24wk, indicating a progressive decline in mitochondrial function with advancing LV dysfunction (Fig. S6A-C). In parallel, pro-fibrotic pathways were positively enriched in Foz+WD 12wk compared to WT+NC 24wk and in Foz+WD 24wk compared to Foz+WD 12wk, consistent with a stepwise worsening of fibrotic remodeling (Fig. S6A-C).

Consistent with transcriptomic evidence of impaired mitochondrial bioenergetics, protein-level analysis in isolated cardiomyocytes revealed dysregulation of key regulators of calcium handling and mitochondrial function (Fig. 6F). SERCA2a expression was reduced in Foz+WD mice, indicating impaired sarcoplasmic reticulum Ca²⁺ reuptake. EMRE, a core component of the mitochondrial calcium uniporter complex, was transiently increased at 12wk, suggesting an early compensatory adaptation, but declined by 24wk in Foz+WD cardiomyocytes. In contrast, MCUb, a negative regulator of mitochondrial calcium uptake, was markedly increased in Foz+WD 24wk hearts (Fig. 6F). Together, these alterations indicate progressive disruption of excitation-contraction coupling and mitochondrial calcium handling during disease progression.

Overall, these cardiac analyses indicate that the progression from moderate to severe LV dysfunction is associated with a sustained reduction in fatty acid oxidation, impaired mitochondrial bioenergetics, and increasing pro-fibrotic signaling, ultimately contributing to the development of HFpEF-like cardiac dysfunction and reduced survival.

### 3.8 LV gene expression and mitochondrial ultrastructural changes during disease regression

The impact of lifestyle modification on cardiac transcriptional remodeling remains poorly understood. To examine this, we compared LV transcriptomes from the regression group with three experimental cohorts: WT+NC 24wk, Foz+WD 12wk, and Foz+WD 24wk (Fig. 7A). Cardiac gene expression in regression mice was first compared with Foz+WD 24wk and then plotted alongside expression changes from Foz+WD 12wk versus Foz+WD 24wk, enabling simultaneous visualization of molecular trajectories during both disease progression and regression (Fig. 7B). Genes consistently upregulated in Foz+WD 24wk hearts across both comparisons, designated as heart failure (HF)-associated genes, were enriched in inflammatory, hypoxia-responsive, and epithelial-to-mesenchymal transition (EMT) pathways (Fig. 7B-C), consistent with ongoing fibrotic remodeling and microvascular rarefaction characteristic of human HFpEF[34]. In contrast, genes selectively upregulated in the regression group were enriched for mitochondrial respiration, ATP synthesis, sarcomere organization, and cardiac muscle differentiation and contraction (Fig. 7B-C). Functional annotation of these regression-specific genes indicates activation of a reparative program characterized by re-engagement of the fetal gene network, a conserved adaptive response that promotes myocardial regeneration and recovery[35]. GSEA comparing Foz+WD 24wk and regression hearts further validated these findings (Fig. S7A). Cellular deconvolution analysis showed that dietary intervention mitigated the expansion of fibroblast and myeloid populations (Fig. S7B).

**Fig. 7:**
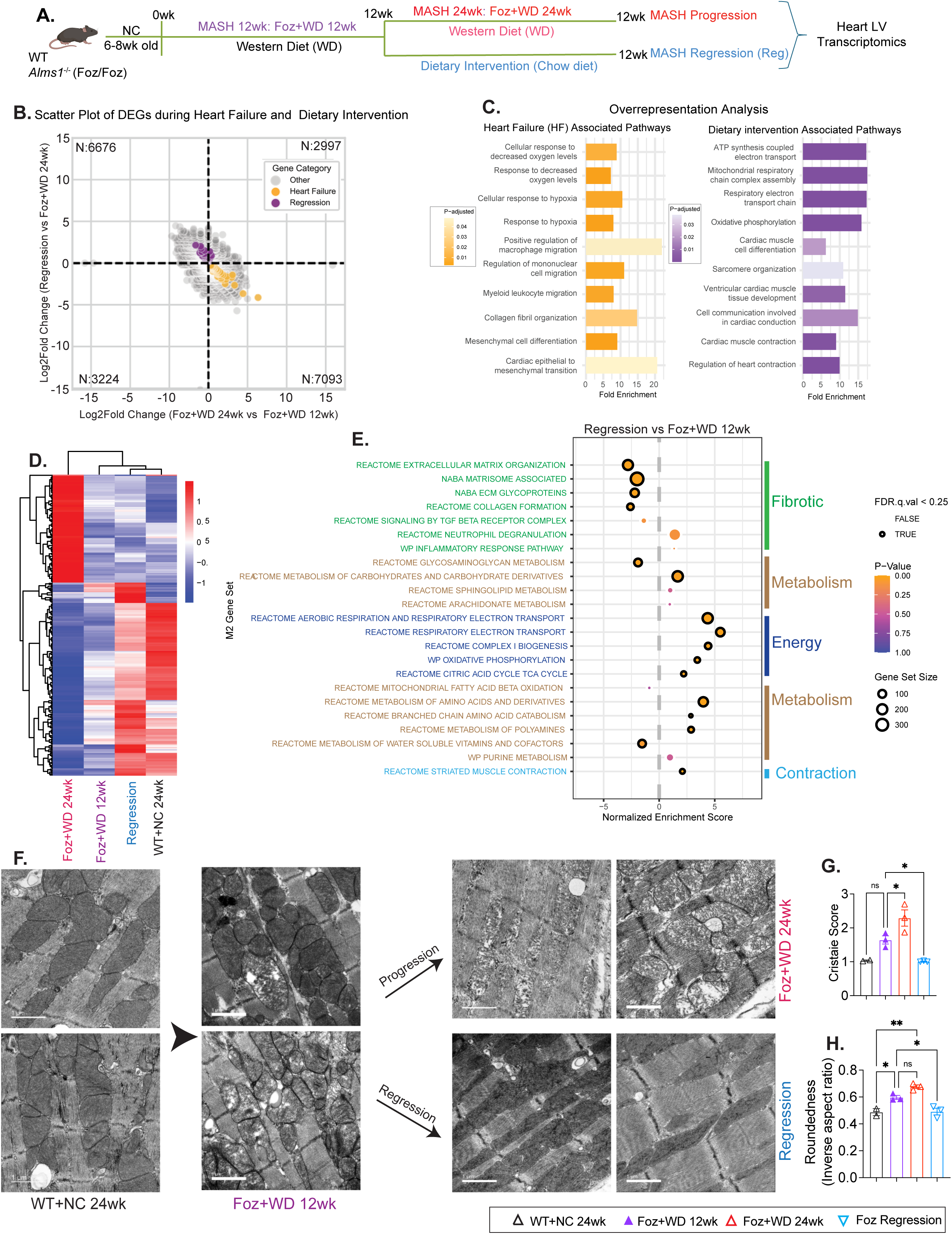
Dynamic changes in LV gene expression and mitochondrial ultrastructure during disease regression. (A) Study design: Foz/Foz mice were fed Western diet (WD) to induce MASH-fibrosis over 12 to 24wk. A subset of 12wk MASH-fibrosis mice were switched to a normal chow diet for 12 additional weeks (MASH and liver fibrosis regression). (B) Scatter plot comparing gene expression changes between two comparisons: Foz+WD 24wk vs. Foz+WD 12wk (x-axis) and Regression vs. Foz+WD 24wk (y-axis). Genes consistently upregulated in Foz+WD 24wk compared to both Foz+WD 12wk and regression are highlighted in yellow (91 genes) and are the gene signature associated with heart failure. The genes uniquely upregulated in the regression model compared to both Foz+WD 12wk, 24wk are highlighted in violet (162 genes) and are the gene signature associated with dietary intervention. (C) Functional enrichment analysis of genes upregulated during heart failure (left; yellow bars) and during dietary intervention (regression, right; violet bars). Pathways are plotted with fold enrichment values on the x-axis, while the color scale represents the adjusted p-value, indicating statistical significance. (D) Heatmap of differentially expressed genes (DEGs) in LV tissue showing consistent transcriptional changes between severe HFpEF-like cardiac dysfunction (Foz+WD 24wk, leftmost group) and healthy controls (WT+NC 24wk, rightmost group). Genes are hierarchically clustered, and z-score normalized, revealing coordinated upregulation (red) and downregulation (blue) patterns on average for n=3 samples in each group. Intermediate-stage HFpEF-like cardiac dysfunction (Foz+WD 12wk) and regression (WD→chow) groups occupy central positions, with the latter clustering closer to healthy controls. This pattern indicates partial restoration of the cardiac transcriptome and suggests reversibility of HFpEF-associated molecular remodeling following dietary intervention (see also figure S7D). (E) Scatter plot illustrating gene set enrichment analysis (GSEA) using the M2 Curated gene set to compare regression vs Foz+WD 12wk. The size of each dot represents the gene set size, dark circles indicate significant enrichment based on FDR q-val<0.25, and the color corresponds to the p-value. (F) Representative TEM micrographs of LV myocardium with corresponding quantifications and scoring criteria (right panel), showing progressive mitochondrial damage associated with worsening fibrosis and HFpEF-like cardiac dysfunction in Foz/Foz mice, and restoration of ultrastructural integrity following fibrosis resolution in the dietary intervention (regression) group. Representative images from two different mice/group are shown. Scale bar=1μm. (G, H) TEM Quantifications: Cristae Score (G): Semi-quantitative assessment of mitochondrial inner membrane integrity using an established scoring system (Flameng et. al., 1980; Vu et. al., 2023). Scores were assigned based on the proportion of mitochondrial area lacking defined cristae architecture (score 1: normal; score 2: ∼25%; score 3: ∼50%; score 4: ∼75%), with increasing scores indicating progressive cristae disorganization. Mitochondrial Roundness (H): Quantification of mitochondrial swelling and loss of elongation. Roundness was calculated by the inverse aspect ratio (minor axis/major axis) (Kalkhoran et. al. 2017) as described in the methods. A higher roundness value (approaching 1.0) indicates a perfectly spherical, swollen organelle. Data are presented as mean ± SEM. Individual data points (triangles) represent independent biological replicates per group (n=2 for WT+NC; n=3 for the remaining groups). At least five independent fields of view (10,000x magnification) were acquired per animal, and a minimum of 70-100 randomly selected, non-overlapping mitochondria were analyzed per sample for both cristae scoring and morphometric calculations. Values were averaged per animal to generate a single biological replicate. Statistical significance was determined by a One-Way ANOVA followed by Sidak’s multiple comparisons test. *p < 0.05, **p < 0.01; ns, not significant.

Leading-edge analysis identified genes driving the observed pathway shifts (Fig. S7C-D). The regression group’s transcriptomic profile closely resembled that of healthy WT+NC controls, indicating broad restoration of cardiac gene expression (Fig. S7C-D). In contrast, Foz+WD 12wk mice showed early dysfunction signatures resembling Foz+WD 24wk hearts (Fig. S7C-D). Key genes mediating cardiac contraction, including troponins (*Tnni3, Tnnt1*) and myosins (*Myl2, Myl3*), were upregulated in regression hearts (Fig. S7C-D), consistent with improved LV contractile reserve and relaxation after dietary intervention (Fig. 5J-L).

Hierarchical clustering confirmed that regression profiles aligned with WT+NC controls, while Foz+WD 24wk remained most divergent and Foz+WD 12wk intermediate (Fig. 7D, Fig. S7E). GSEA comparing regression to Foz+WD 12wk revealed enrichment of mitochondrial bioenergetic and contractile pathways, with suppression of pro-fibrotic and disease-associated signaling (Fig. 7E). These data demonstrate that dietary intervention improves pathological transcriptional remodeling and promotes functional recovery.

Transmission electron microscopy (TEM) revealed pronounced ultrastructural remodeling and sarcomeric disorganization in Foz+WD hearts compared with healthy controls (Fig. 7F-H, Fig. S7F). Cardiomyocytes displayed disrupted cristae (Fig. 7G) [36, 37] and swollen mitochondria (Fig. 7H)[38], hallmarks of metabolic and contractile stress that worsened with disease progression. Remarkably, these structural abnormalities were largely reversed following dietary intervention, consistent with transcriptomic signatures of restored mitochondrial metabolism and contractile function in fibrosis-regressed mice (Fig. 7F-H, Fig. S7F).

## 4. Discussion

### 4.1 Modeling the Liver-Heart Axis in Metabolic Disease

The intersection of MASLD and cardiovascular disease, particularly HFpEF, represents a growing clinical challenge with limited mechanistic insight. Although these conditions frequently coexist, the molecular and physiological links between hepatic and cardiac pathology remain poorly defined[39]. Progress has been hindered by the lack of translational preclinical models that recapitulate the complex, multi-organ phenotype observed in patients[15, 40]. While several rodent models, including the Golden Syrian hamster[40] and the stroke-prone spontaneously hypertensive rat (SHRSP5/Dmcr)[41], have been used to study MASH associated HFpEF, none recapitulate mortality, a defining feature of human HFpEF. Moreover, C57BL/6N mice fed with high fat, high fructose, high cholesterol diet, do not develop HFpEF despite developing MASH with moderate fibrosis[42]. Here, we address this gap using the Foz+WD mouse model, which recapitulates the cardinal features of human cardiometabolic dysfunction, including MASH with advanced fibrosis[17], HCC[17, 43, 44], systemic insulin resistance, renal impairment[20], cardiac contractile abnormalities, and premature mortality.

Our findings also support a reversible relationship between liver fibrosis and cardiac dysfunction. Male Foz+WD mice developed metabolic syndrome and progressive hepatic injury, culminating in advanced liver fibrosis and high mortality by 34wk, closely mirroring human MASLD, where liver fibrosis stage is a key determinant of mortality risk[45, 46]. Importantly, worsening hepatic pathology coincided with the onset of cardiac dysfunction exhibiting key features of HFpEF such as concentric hypertrophy, ventricular dilation, and slightly reduced ejection fraction (∼57%)[47], accompanied by impaired β-adrenergic responsiveness, diminished contractile reserve, and elevated plasma BNP. In contrast, control groups with MetS or MASH but without liver fibrosis did not develop cardiac dysfunction, suggesting that advanced hepatic injury strongly correlates with cardiac remodeling.

### 4.2 Mitochondrial Remodeling and Mechanistic Insights

The transcriptomic, functional, and ultrastructural analyses collectively suggest that cardiac remodeling in Foz+WD mice is not simply a passive consequence of systemic metabolic stress, but reflects an active and coordinated myocardial state transition. The combined suppression of fatty-acid β-oxidation, amino-acid catabolism, and oxidative phosphorylation, together with induction of inflammatory and matrix-remodeling programs, is consistent with a heart that has shifted from a high-efficiency oxidative organ to a stress-adapted, energetically constrained phenotype. Mitochondrial injury represents an important feature linking metabolic overload, impaired contractile reserve, and structural remodeling, thereby linking molecular remodeling to functional HFpEF-like physiology. The ultrastructural evidence of swollen mitochondria and cristae disruption further suggests that the defect lies not only in substrate selection but in the capacity of the myocardium to convert nutrient availability into usable ATP.

Importantly, the substantial reversal of these changes with dietary normalization indicates that this cardiac phenotype retains substantial biological plasticity, rather than representing fixed end-stage damage, highlighting the therapeutic potential of timely intervention along the metabolic-HFpEF axis.

Comparisons between the Foz+WD cardiac transcriptome and failing human HFpEF myocardium revealed conserved signatures of immune activation, mitochondrial dysfunction, and structural remodeling, including hypoxia-inducible and microvascular rarefaction pathways[34, 48, 49]. Together, these data identify mitochondrial dysfunction and impaired bioenergetics as key features of MASLD-associated cardiac pathology and underscore the model’s translational relevance[49–52].

### 4.3 Metabolic Restoration and Cardiac Recovery

The Foz+WD model also demonstrated the reversibility of the liver-heart axis through dietary modification. Switching Foz+WD mice to chow after 12wk, when MASH and moderate fibrosis were already established, reversed hepatic steatosis and liver fibrosis, with parallel improvement in cardiac performance. Regression mice showed restored contractility, improved dobutamine responsiveness, lower plasma BNP, and enhanced sarcomere shortening, reflecting substantial functional recovery. Although cardiac hypertrophy persisted, these improvements prevented progression to severe HFpEF-like cardiac dysfunction and improved survival from ∼20% to 100%, emphasizing the cardioprotective benefits of metabolic normalization.

LV transcriptomic profiling during regression revealed reactivation of reparative and energy-restorative programs. Genes involved in mitochondrial respiration, ATP synthesis, sarcomere organization, and muscle differentiation were upregulated, while fibrosis and disease-associated pathways were suppressed. The LV transcriptional profile of regression mice closely resembled that of healthy controls, with increased expression of contraction genes, consistent with functional recovery. Electron microscopy corroborated these findings, showing reversal of mitochondrial and sarcomeric disorganization. These molecular and structural changes paralleled the physiological recovery observed with dietary intervention and highlight the heart’s intrinsic capacity for repair once systemic metabolic stress is reduced.

Our study provides important insights into the molecular pathways in the LV that are disrupted during metabolic dysfunction associated HFpEF and those that support recovery during disease regression. The Foz+WD model provides a translational platform for probing putative liver-heart crosstalk and evaluating interventions targeting metabolic and mitochondrial pathways. Importantly, our results demonstrate that timely dietary intervention, a cornerstone of MASLD management, can reverse hepatic fibrosis, halt cardiac decline, restore myocardial function, and markedly improve survival. These findings underscore the need for integrated strategies that address liver and cardiac pathology together to optimize clinical outcomes[53–56].

### 4.4 Strengths and Limitations

The Foz/Foz model provides a robust and highly penetrant model that recapitulates the full spectrum of MASLD progression, including advanced fibrosis, cirrhosis, HCC, CKD, and HFpEF-like cardiac dysfunction, with strong translational and multi-organ relevance. Its responsiveness to dietary intervention further enable interrogation of disease plasticity and therapeutic mechanisms, including regression of established pathology.

Beyond hyperphagia, whether loss of Alms1 exerts direct effects on cardiomyocytes remains to be determined, although Foz/Foz mice maintained on a normal chow diet do not develop heart failure within the time frame examined. Notably, the HFpEF phenotype is reversible with dietary intervention. Interestingly, Foz/Foz mice on a NOD.B10 background fed a high-fat diet (HFD) exhibited minimal cardiac dysfunction and did not progress beyond cardiac hypertrophy[57]. This contrasts with our studies in Foz/Foz mice on a C57BL/6 background fed a WD containing 0.2% cholesterol, highlighting the importance of genetic background and diet composition in driving cardiac disease severity. Also, attribution of mortality to cardiac causes are not definitive despite its strong association with impaired β-adrenergic reserve. In accordance, our data did not indicate major hepatic or renal contributions to mortality.

The observed cardiac phenotype captures multiple features consistent with HFpEF, including preserved ejection fraction (>50%), LV hypertrophy, impaired contractility, reduced β-adrenergic reserve, elevated BNP, and increased relaxation constant (ρ). However, additional assessments, including pressure-volume analysis, direct filling pressures, pulmonary hemodynamics, exercise testing and direct assessment of mitochondrial respiration were not performed and would further refine phenotypic classification. Additionally, HFpEF in humans is defined by integrated clinical criteria that cannot be fully recapitulated in mice and, therefore, it is best interpreted as HFpEF-like phenotype rather than a definitive clinical entity.

Cross-species comparisons with human HFpEF datasets were limited due to the lack of well-defined human HFpEF datasets as well as heterogeneity of underlying HFpEF causality. In addition, cardiomyocyte contractility studies were performed with relatively small group sizes (n=3 or 4 mice/group), and these findings should therefore be interpreted cautiously despite the consistency across independent functional readouts.

While this study provides comprehensive transcriptional and functional characterization of cardiac dysfunction, alongside prior hepatic[17, 19]and renal[20] analyses in this model, the mechanisms underlying inter-organ crosstalk require further investigation and the specific mediators of liver-heart communication are yet to be identified[58]. Although liver fibrosis that develops during MASH is strongly associated with cardiac dysfunction, the study design does not allow causal inference, as metabolic, hepatic, and cardiac abnormalities evolve in parallel during both progression and regression. Metabolic comorbidities, including hyperglycemia and insulin resistance, may independently contribute to mitochondrial dysfunction and cardiac remodeling and cannot be fully disentangled. Further studies are required to define the causal contributions of MASH and liver fibrosis to HFpEF, and vice-versa.

Despite these limitations, this model provides a translationally relevant platform for investigating putative liver-heart crosstalk and, more broadly, recapitulates the integrated cardio-renal-hepatic manifestations of metabolic dysfunction, enabling future interrogation of mechanisms associated with multi-organ disease progression in MASLD.

## Acknowledgments

Research was supported by NIH Grants to D.D. (R01DK133930), the San Diego Digestive Diseases Research Center (SDDRC) (NIH DK120515) and Sanford Burnham Prebys Medical Discovery institute (SBP) NCI Cancer Center Support grant P30CA030199 for the core services. The authors would like to thank the University of California, San Diego (UCSD) Cellular and Molecular Medicine Electron Microscopy Core (UCSD-CMM-EM Core, RRID:SCR_022039) for TEM sample preparation and microscope access, Seaweed cardiovascular physiology core services for hemodynamics and echocardiogram studies; UCSD and SBP histopathology core facility for tissue processing and SBP bioinformatics core for computational analysis. We also thank Dr. Xin Tu (Professor of Biostatistics, Division of Biostatistics and Bioinformatics, UCSD) for help with statistics. Graphical abstract was created with BioRender.com.

## Author contributions

S.G., T.K., D.A.B., and D.D. designed research; S.G., B.G., Y.G., J.S., G.G, K.I., R.M., performed experiments and analyzed data; W.D., K.P., E.A., analyzed data and provided feedback; D.D. conceptualized and supervised research; and S.G., B.G., D.A.B., and D.D. wrote the paper.

