## Supplementary Materials for "A Translational Model of MASLD-Associated HFpEF Defines Mitochondrial Dysfunction and Cardiac Plasticity During Disease Progression and Regression"

### SUPPLEMENTARY METHODS

#### Isolation of Cardiac Tissue

Cardiac tissue was collected following diastolic arrest unless otherwise noted. Briefly, mice were anesthetized with isoflurane, after which 0.3 M KCl was injected into the heart to induce rapid diastolic arrest. The heart was then excised and washed in 0.3 M KCl to remove residual blood. For experiments requiring left ventricular (LV) tissue, the arrested heart was transferred to ice-cold phosphate-buffered saline (PBS), and the LV was identified and carefully dissected away from the right ventricle and interventricular septum. LV tissue was immediately processed for downstream applications.

#### Histology and Image Quantification

Tissue samples, including the left lateral liver lobe and whole heart, were fixed in 10% neutral-buffered formalin for 48 hours, followed by paraffin embedding and sectioning. Liver sections were stained with hematoxylin and eosin (H&E) and Picro Sirius Red to assess general morphology and collagen deposition, respectively. Cardiac sections were stained with Wheat Germ Agglutinin (WGA; RL-1022, Vector Laboratories) for delineation of cardiomyocyte size and immunostained for perilipin 2 (NB110-40877SS, Bio-Techne) to evaluate lipid accumulation. For each tissue type, seven non-overlapping fields per section were randomly imaged using an Olympus microscope. Quantitative analysis was performed using ImageJ (NIH), and Sirius Red and immunohistochemical signals were normalized to the non-lipid area determined from H&E-stained sections to control for the extent of steatosis.

#### RNA sequencing

The paired-end reads that passed Illumina filters were filtered for reads aligning to tRNA, rRNA, adapter sequences, and spike-in controls. The reads were then aligned to the GRCm38.p4 reference genome and Gencode M9 annotations using STAR (v 2.6.1)[1]. DUST scores were calculated with PRINSEQ Lite (v 0.20.3) [2] and low-complexity reads (DUST > 4) were removed from the BAM files. The alignment results were parsed via the SAMtools [3] to generate SAM files. Read counts to each genomic feature were obtained with featureCounts (v 1.6.5) [4]. After removing absent features (zero counts in all samples), the raw counts were then imported to the R/Bioconductor package DESeq2 (v 1.24.0) [5] to identify differentially expressed genes among samples. P-values for differential expression are calculated using the Wald test for differences between the base means of two conditions. These P-values are then adjusted for multiple test correction using the Benjamini-Hochberg algorithm. Genes that are consistently upregulated and downregulated were identified using linear regression. Gene set enrichment analysis was done using the 'GseaPreranked' method with the 'classic' scoring scheme at GSEA Software (v4.4.0). Rank files for each DE comparison of interest were generated by calculating $\pi$-value = $-\log10(\text{p-adj}) \times \text{LFC}$ [6].

**Deconvolution analysis**

Processed snRNA-seq datasets from mouse LV (control condition, n = 4) were obtained from GEO (GSE263648)[7]. Data were reanalyzed in R (v4.5.2) using Seurat (v5.4.0). Doublets were identified using scds (v1.26.1) and, together with cells exhibiting >10% mitochondrial content, were excluded. Data were normalized using NormalizeData, and the top 2,000 highly variable genes were identified using FindVariableFeatures. Scaling was performed using ScaleData with regression of nCount_RNA and mitochondrial percentage. Principal component analysis was conducted using RunPCA on highly variable genes.

Samples were integrated using IntegrateLayers with Harmony-based integration (method = "HarmonyIntegration", orig.reduction = "pca"). Dimensionality reduction and clustering were performed using RunUMAP, FindNeighbors, and FindClusters (resolution = 0.5), with principal components explaining ~90% of variance. Cluster marker genes were identified using FindAllMarkers, and cell types were annotated based on established marker gene expression. Residual doublets identified by aberrant marker expression were excluded from downstream analyses.

To construct a reference signature matrix, the integrated dataset was downsampled to a maximum of 500 cells per cell type and restricted to the top 10,000 highly variable genes. The normalized expression matrix was used as input for the CIBERSORTx "Create Signature Matrix" module to generate a gene signature matrix. Cell type proportions in bulk RNA-seq samples were then estimated using the CIBERSORTx "Impute Cell Fractions" module.

**Transmission Electron microscopy**

LV samples collected as described above were prepared for transmission electron microscopy (TEM). Immediately following dissection, LV tissue fragments (~1-2 $mm^3$) were fixed with 2% paraformaldehyde + 2.5% glutaraldehyde in 0.15 M sodium cacodylate buffer and further postfixed in 1% $OsO_4$ in 0.15 M cacodylate buffer for 1hr on ice. The specimens were stained with 2% uranyl acetate for 1hr on ice, following graded dehydration in series of ethanol (50-100%) while remaining on ice. The cells were then subjected to one wash with 100% ethanol and two washes with acetone (15min each) and embedded with Durcupan (Sigma-Aldrich #44610-1EA). Sections were cut at 60 nm on a Leica UCT ultramicrotome and picked up on 300 mesh copper grids. Sections were post-stained with 2% uranyl acetate for 5 minutes and Sato's lead stain (1% lead acetate, 1% Lead Nitrate, 1% Lead citrate and 2% sodium citrate in water) for 1 minute. Finally, the sections were observed at the UCSD Electron Microscopy Core (UCSD-CMM-EM Core, RRID:SCR_022039) using a Jeol 1400 plus operated at 80KeV and equipped with a bottom-mounted Gatan One View camera.

Mitochondrial morphology: Electron micrographs were analyzed using NIH ImageJ software (FIJI, version 2.0.2) to quantify mitochondrial morphology. For each animal (n=3 mice per group), at least five independent fields of view were acquired at 10,000x and 3000x magnifications, and 70-100 intact, non-overlapping mitochondria were randomly selected per mouse for analysis. Mitochondrial shape was assessed by calculating the aspect ratio using the ImageJ Fit Ellipse function, which determines the major and minor axes of the best-fitting ellipse[8]. Roundness was calculated as the inverse aspect ratio (minor axis/major axis), where values approaching 1.0 indicate a spherical, swollen mitochondrial phenotype[9].

Cristae Score: Cristae integrity was evaluated using a semi-quantitative scoring system[10, 11]. Scores were assigned based on the proportion of mitochondrial area lacking defined cristae architecture (score 1: normal; score 2: ~25%; score 3: ~50%; score 4: ~75%), with increasing scores indicating progressive cristae disorganization. To avoid pseudoreplication, all morphometric and scoring measurements were averaged per animal to generate a single biological replicate (n = 3 per group except healthy control WT+NC which is n=2), which was used for all statistical analyses.

**Transthoracic Echocardiography**

Before echocardiography, a depilatory cream is applied to the anterior chest wall to remove hair. Mice are anesthetized with 5% isoflurane for 15 seconds, then maintained at 0.5% isoflurane throughout the echocardiographic examination. Small needle electrodes are inserted into one upper and one lower limb for simultaneous electrocardiogram (ECG) recording. Transthoracic echocardiography, including M-mode and 2-dimensional imaging, is performed using the FUJIFILM VisualSonics Inc. Vevo 2100 high-resolution ultrasound system with a linear transducer (32-55 MHz). Measurements of chamber dimensions and wall thickness are taken. Percentage fractional shortening (%FS) is used as an indicator of systolic cardiac function. Pulsed wave Doppler in the apical 4-chamber view is employed to acquire the ratio of peak velocities of early to late mitral inflow (E/A) and deceleration time (DT). Additionally, tissue Doppler imaging in the apical 4-chamber view allows for the measurement of mitral annular motion velocities, with E' and A' representing the peak mitral annular velocities during early and late filling, respectively. To ensure proper E and A wave separation, additional ketamine (50 mg/kg) is administered.

**Quantitative Real-Time PCR (qRT-PCR)**

Total RNA was extracted using RNeasy Mini columns (Qiagen, Valencia, CA), and reverse transcription was performed according to standard protocols. qRT-PCR was conducted using the QuantStudio 5 system (Applied Biosystems, Carlsbad, CA), and gene expression levels were normalized to *Hprt* using the ΔΔCt method. Primers were designed via PrimerBank (https://pga.mgh.harvard.edu/primerbank/). Data are presented either as fold change relative to the indicated control or as expression normalized to a housekeeping gene.

Following primers (mouse) were used:

| Gene | Forward primer | Reverse primer |
|---|---|---|
| *Cd11b* | CCATGACCTTCCAAGAGAATGC | ACCGGCTTGTGCTGTAGTC |
| *Hprt* | GTTAAGCAGTACAGCCCCAAA | AGGGCATATCCAACAACAAACTT |
| *F4/80* | ACCAGAGGAAATTTTCAATAGGC | TGATGCACTTGCAGAAAACA |
| Ly6g | CCTGCAACACAACTACCTGCCCC | AGTGGGGCAGATGGGAAGGC |

**Western Blot**

Cardiomyocytes isolated as described before were homogenized in lysis buffer (20 mM Tris (pH 7.4), 20 mM NaCl, 0.1 mM EDTA, 0.025 mM PUGNAc, 1% Triton X- 100, 1:1000 diluted protease, and phosphatase inhibitor mixture). 50-100 µg protein samples were loaded on NuPAGE 4-12% Bis-Tris gels (Invitrogen). Separated proteins were transferred to PVDF membranes. anti-SERCA2 (sc-376235), anti-EMRE (Santa Cruz SC-86337), and anti-MCUb (“CCDC109B” produced by ProSci Inc.) were used as primary antibodies. Anti-mouse IgG-horseradish peroxidase (HRP)-conjugated (ThermoFisher Scientific 31430) and anti-Rabbit IgG-HRP-conjugated (Cell Signaling 7074) were used as secondary antibodies. Total protein stain (Thermo Scientific 24585) was used as loading standards. Band density was quantified with ImageJ software as previously described.

**Metabolic Phenotyping**

Whole-body energy homeostasis was assessed using the Comprehensive Lab Animal Monitoring System (CLAMS; Columbus Instruments, OH). 6-months WD fed Male Foz/Foz (n=4) and WT (n=5) mice assessed for metabolic phenotyping. Mice were individually housed in metabolic chambers with ad libitum access to WD and water. After a 24-hour acclimation, continuous metabolic data were recorded over 48 hours under standard light/dark conditions. Parameters measured included $VO_2$, $VCO_2$, respiratory exchange ratio (RER), energy expenditure (EE), food and water intake, and locomotor activity (horizontal and vertical beam breaks). Data were acquired every 15 minutes and analyzed as light/dark phase-specific and 24-hour averages. EE was calculated using the Lusk equation based on $VO_2$ and $VCO_2$.

**Plasma Biochemistry**

Peripheral blood was collected from the inferior vena cava, and plasma was isolated using BD Microtainer tubes (cat. 365985, BD). ALT, ALP, cholesterol, total bilirubin, albumin, and BUN were measured using the Abaxis Mammalian Liver Profile system (catalog# 500-0040-12). Plasma triglycerides were quantified using the Triglyceride Reagent Set (236-60; Sekisui Diagnostics, PE, Canada). Plasma BNP levels were assessed using a mouse BNP ELISA Kit (NBP2-70011) following manufacturer protocols.

**Cardiac dysfunction staging and correlation with liver/metabolic phenotypes**

To evaluate the association between progressive cardiac dysfunction and metabolic disease severity, mice were assigned to an ordinal cardiac dysfunction stage based on integrated cardiac phenotyping, as summarized in Supplementary Fig. S5C. Stage 0 was defined as healthy control mice. Stage 1 was defined as cardiac hypertrophy alone. Stage 2 was defined as cardiac hypertrophy with cardiomyocyte contractile dysfunction and a mildly impaired β-adrenergic response. Stage 3 was defined as cardiac hypertrophy with cardiomyocyte contractile dysfunction, a severely impaired β-adrenergic response, and increased overall mortality observed in the corresponding disease stage. This staging system was used as an ordinal summary of cardiac disease severity to evaluate its relationship with liver and metabolic parameters.

Associations between cardiac dysfunction stage and liver/metabolic phenotypes (also summarized in Supplementary Fig. S5C)—including liver fibrosis, steatosis, plasma ALT, plasma cholesterol, plasma triglycerides, blood glucose, body weight, and heart weight—were evaluated using Spearman's rank correlation. The relationship between cardiac dysfunction stage and liver fibrosis (Sirius Red–positive area) is shown in a correlation plot, with each point representing one mouse. The plot reports the Spearman correlation coefficient, nominal *P* value, and Benjamini–Hochberg false discovery rate–adjusted *P* value. Associations with the other metabolic parameters are summarized in a Spearman correlation summary plot with Spearman correlation coefficients on the x-axis and statistical significance expressed as −log10(*P* value) on the y-axis.

**Manuscript Preparation**

Some portions of the text were edited for clarity and conciseness using ChatGPT (OpenAI). All content was reviewed and verified for accuracy by the authors.

**Reporting and Study Design Details**

Outcome assessment was performed blinded to group allocation where feasible; histological scoring, morphometric image analyses, plasma biochemistry, and qRT-PCR were conducted using coded samples. Transcriptomics studies were done using 3 mice/group. Sample size (n) or other studies are indicated in the figure legends and in the individual panels as applicable (each dot in bar plots represent one mice). The group sizes represent the number of mice or independent values (not technical replicates). No predefined inclusion or exclusion criteria were applied, and no animals or data points were excluded from analysis. To minimize potential confounders, animals from different groups were processed and measured in parallel and in mixed order whenever feasible; cage location was not treated as an experimental variable. No formal preregistration or publicly registered study protocol was completed prior to the study.

**Supplementary Fig. S1: A pre-clinical model of MetS, MASH and cardiometabolic dysfunction with mortality.** (A) WT+WD 24wk and Foz+WD 24wk mice were placed in instrumented metabolic cages individually (n=5 and 4 respectively) to monitor multiple metabolic parameters as indicated. (B-F) 6-8wk old mice of indicated genotypes were either fed NC or WD for 24wk. Liver and plasma were collected for subsequent analyses. (B) Body weight (BW) and liver weight (LW), across the indicated groups are plotted (Obesity-associated parameters). (C) Plasma fasting blood glucose, plasma cholesterol, plasma triglyceride levels are plotted (biochemical parameters). (D) Circulating alanine aminotransferase (ALT) levels reflecting liver injury. (E) Liver function markers, including total bilirubin, albumin, and blood urea nitrogen (BUN) are plotted. (F) Hepatic mRNA expression of immune cell markers *Cd11b* and *F4/80* (monocytes/macrophages), and *Ly6G* (neutrophils) were assessed by qRT-PCR, normalized to *Hprt* expression and plotted as fold change relative to healthy controls (WT+NC 24wk). Data are presented as mean±SEM; Sample size (n=3-10 mice per group) as indicated in each panel (each dot representing one mouse). Group comparisons were performed using one-way ANOVA (Fig. S1B-F) or two-way ANOVA (Fig. S1A) followed by Sidak's multiple comparisons test. *p<0.05, **p<0.01, ***p<0.001, ****p<0.0001, ns=not significant. (G) Table showing serum clinical chemistry at necropsy in Foz+WD mice (24wk) (n=3).

**Supplementary Fig. S2: Left ventricular dysfunction in MASLD.** (A) Representative Doppler echocardiography images including pulsed-wave (PW) Doppler and tissue Doppler imaging (TDI) used to assess LV filling and myocardial relaxation. (B) Mitral valve early-to-late filling velocity ratio (MV E/A). (C) Mitral valve early filling velocity to early diastolic mitral annular velocity ratio (MV E/E'), an indicator of LV filling pressure. Group comparisons were performed using one-way ANOVA with sample size (n=4-8 mice/group) as indicated in each panel (each dot representing one mouse). As the overall ANOVA was not significant, no post hoc multiple comparisons were performed. (D-E) Hemodynamic assessment at baseline and with increasing dobutamine stimulation: (D) Heart rate (BPM) changes, and (E) Maximum LV pressure ($pressure^{max}$) across indicated groups are plotted along with the slope of the respective curves. Sample size (n) as indicated in Fig. 2K-Q Group comparisons were performed using two-way ANOVA followed by Sidak's multiple comparisons test. Symbols &, #, and $ indicate P < 0.05 for the following comparisons: & : Foz+WD 24wk vs WT+WD 24wk # : Foz+WD 24wk vs Foz+NC 24wk $ : Foz+WD 24wk vs WT+NC 24wk (F) Female Foz+WD 24wk echocardiographic parameters measured and plotted with the other groups (males) for comparison as shown in Figure 2B, I, and J. One-way

ANOVA followed by Sidak's multiple comparisons test when applicable. **p < 0.01, ns=not significant. All data in the figure are presented as mean ± SEM.

**Supplementary Fig. S3: LV Transcriptomics reveals key pathways associated with LV dysfunction.** (A) Heatmap showing the relative expression of genes from analysis in Fig. 3C and 3D. (B-C) Scatter plot illustrating GSEA between (B) Foz+WD 24wk vs WT+WD 24wk and (C) Foz+WD 24wk vs Foz+NC 24wk, using the M2 Curated gene set. The size of each dot represents the gene set size, dark circles indicate significant enrichment based on FDR q-val < 0.25, and the color corresponds to the p-value. (D) Representative Sirius red (SR) stained mouse heart sections of the indicated groups. (E-F) UMAP visualization of re-clustered mouse LV snRNA-seq data obtained from Li et al. (GSE263648) (E). Cell type identities were assigned using expression of canonical marker genes shown in the dotplot (F).

**Supplementary Fig. S4: Effect of MASH-fibrosis regression on cardiac remodeling.** (A) Body weight and liver weight measurements in the indicated groups. (B-E) Plasma metabolic and liver injury markers were measured. (B) Triglycerides; (C) cholesterol; (D) alanine aminotransferase (ALT); (E) alkaline phosphatase (ALP). (F, G) M-mode echocardiographic measurements of LV: (F) ejection fraction (EF); (G) interventricular septum thickness at diastole (IVSd), posterior wall thickness at diastole (LVPWd), LV internal diameter at diastole (LVIDd), internal diameter at systole (LVIDs), and posterior wall thickness at systole (LVPWs). (H) Representative Doppler echocardiographic images, including pulsed-wave (PW) Doppler and tissue Doppler imaging (TDI), used to assess LV filling and myocardial relaxation. (I-J) Diastolic function indices were measured and plotted: (I) Mitral valve early (E) to atrial (A) wave velocity ratio (MV E/A) and (J) Mitral valve early (E) to early mitral annular velocity (E') ratio (MV E/E'). Data are presented as mean ± SEM. Sample size (n=4-6 mice/group), as indicated in each panel (each dot representing one mouse). Group comparisons were performed using one-way ANOVA followed by Sidak's multiple comparisons test when applicable. *p<0.05, **p<0.01, ***p<0.001, ****p<0.0001, ns=not significant.

**Supplementary Fig. S5: Dietary intervention improves LV dysfunction.** (A-B) Hemodynamic parameters under dobutamine stress: (A) maximum left ventricular pressure ($Pressure^{max}$); (B) heart rate (beats per minute, BPM), along with corresponding slopes of response curves across the indicated groups. Group comparisons were performed using two-way ANOVA followed by Sidak's multiple comparisons test. Symbols, &, # and $ indicate P < 0.05 for the respective comparisons. & : Foz+WD 24wk vs WT+NC 24wk, # : Foz+WD 24wk vs

Foz+WD 12wk, $ : Foz+WD 24wk vs Regression 12wk (C) Summary table showing metabolic, hepatic, and cardiac phenotypes across different experimental groups.

**Supplementary Fig. S6: LV remodeling pathways and gene expression changes during MASH progression.** (A-C) Bubble scatter plot illustrating gene set enrichment analysis (GSEA) using the M2 Curated gene set to compare (A) Foz+WD 24wk vs WT+NC 24wk, (B) Foz+WD 24wk vs Foz+WD 12wk (C) Foz+WD 12wk vs WT+NC 24wk. The size of each dot represents the gene set size, dark circles indicate significant enrichment based on FDR q-val< 0.25, and the color corresponds to the p-value.

**Supplementary Fig. S7: LV gene expression and mitochondrial ultrastructural changes during disease regression**. (A) GSEA comparing Foz+WD 24wk vs regression groups using the M2 curated gene set. Each dot represents an individual gene set, with size indicating the number of genes, color representing P value, and dark circles denoting significantly enriched pathways (FDR q < 0.25). (B) Estimated cell-type proportions derived from bulk cardiac RNA-seq cell deconvolution using CIBERSORTx. Shown are fibroblast, myeloid cell and cardiomyocyte fractions across groups. Group comparisons were performed using pairwise t-test with Benjamini-Hochberg (BH) correction, q values are indicated. (C-D) Heatmap showing the genes (Z-score) driving (C) the heart failure associated and (D) regression-associated pathways (related to Figure 7B, C). (E) Cosine similarity index (%) (based on Figure 7D) of LV transcriptomes from Foz+WD 24wk, Foz+WD 12wk, and regression (WD to chow) groups compared to healthy WT+NC 24wk controls. Foz+WD 24wk LV showed the lowest similarity, reflecting advanced disease-associated transcriptional reprogramming. In contrast, Foz+WD 12wk mice exhibited intermediate similarity, and regression mice displayed the highest similarity to healthy transcriptomes, indicating substantial molecular restoration following dietary intervention. (F) Representative TEM images from two different mouse/group are shown (lower magnification of Fig. 7F). Scale bar=2μm.

Supplementary Figure S1

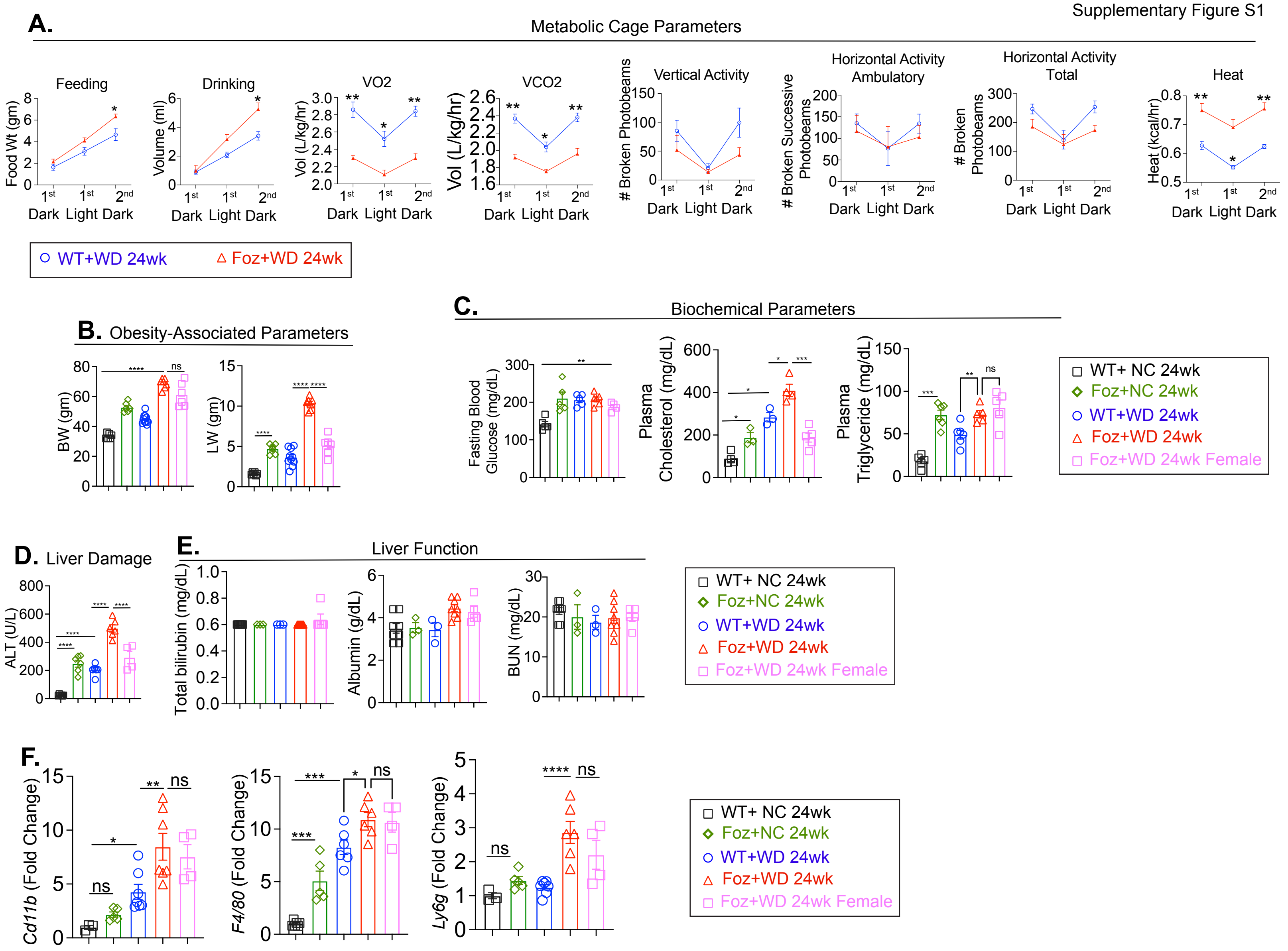


**G.**

| | | Sample ID | FOZ+WD 24wk # 1 | FOZ+WD 24wk # 2 | FOZ+WD 24wk # 3 |
|---|---|---|---|---|---|
| **Analyte name** | **Units** | **Reference ranges** | | | |
| Albumin | G/dl | 2.5 - 4.8 | 4.7 | 4.7 | 4.7 |
| Alkaline phosphatase | U/l | 62 - 209 | 280 | 252 | 247 |
| Alanine transaminase | U/l | 28 - 132 | 846 | 759 | 757 |
| Amylase | U/l | 1691 - 3615 | 848 | 1013 | 864 |
| Blood urea nitrogen | Mg/dl | 18 - 29 | 22 | 17 | 36 |
| Calcium | Mg/dl | 5.9 - 9.4 | 11.6 | 11.3 | 11.9 |
| Phosphorus | Mg/dl | 6.1 - 10.1 | 7.9 | 6.5 | 9.4 |
| Creatinine | Mg/dl | 0.2 - 0.8 | 0.4 | 0.2 | 0.3 |
| Sodium | Mmol/L | 126 - 182 | 158 | 153 | 157 |
| Potassium | Mmol/L | 4.7 - 6.4 | 7 | 6.2 | Hem |
| Total protein | G/dl | 3.6 - 6.6 | 8 | 7.8 | 4.6 |
| Globulin | G/dl | N/a | 3.4 | 3.1 | 1.4 |

Supplementary Figure S2

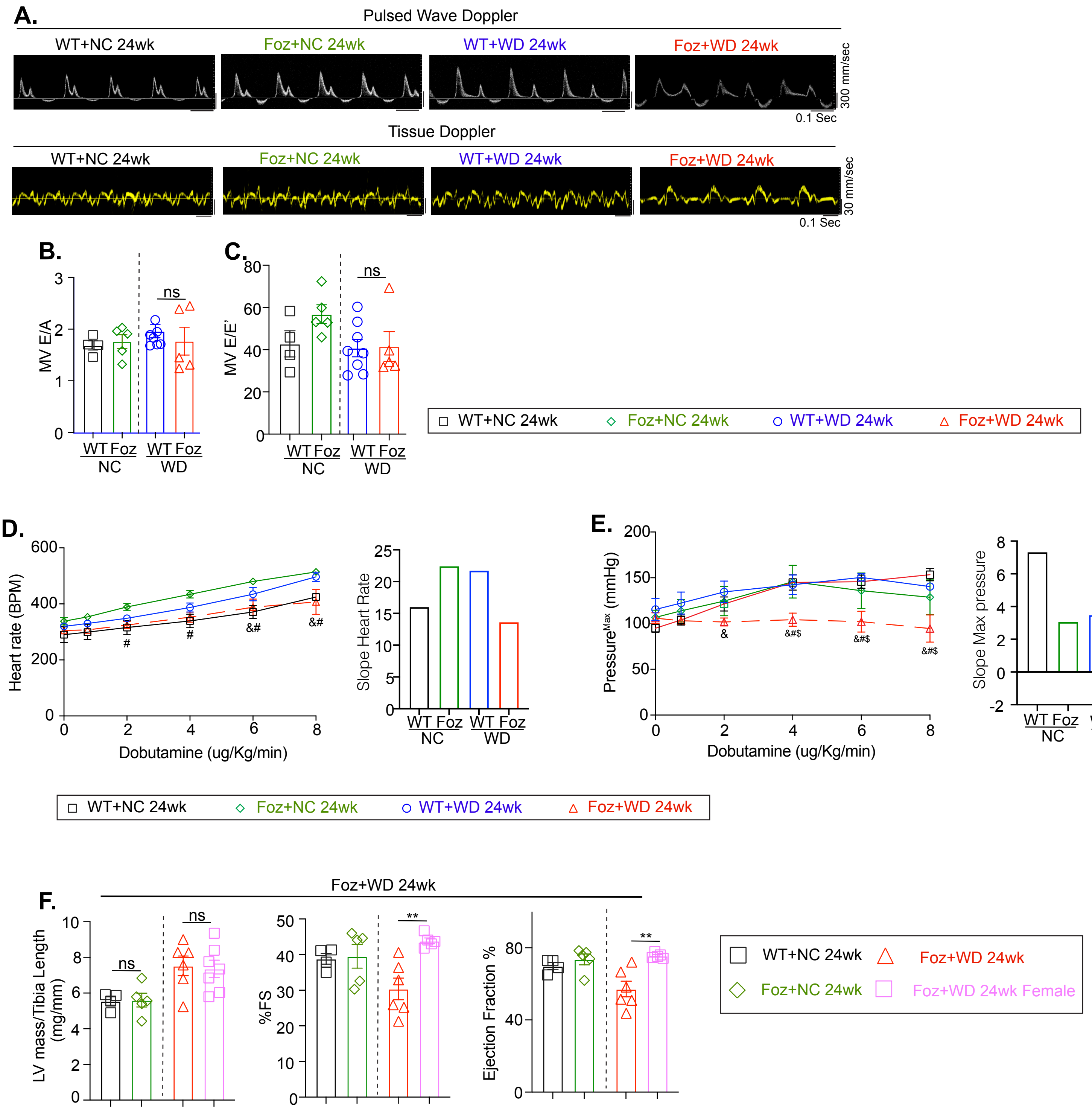

Supplementary Figure S3

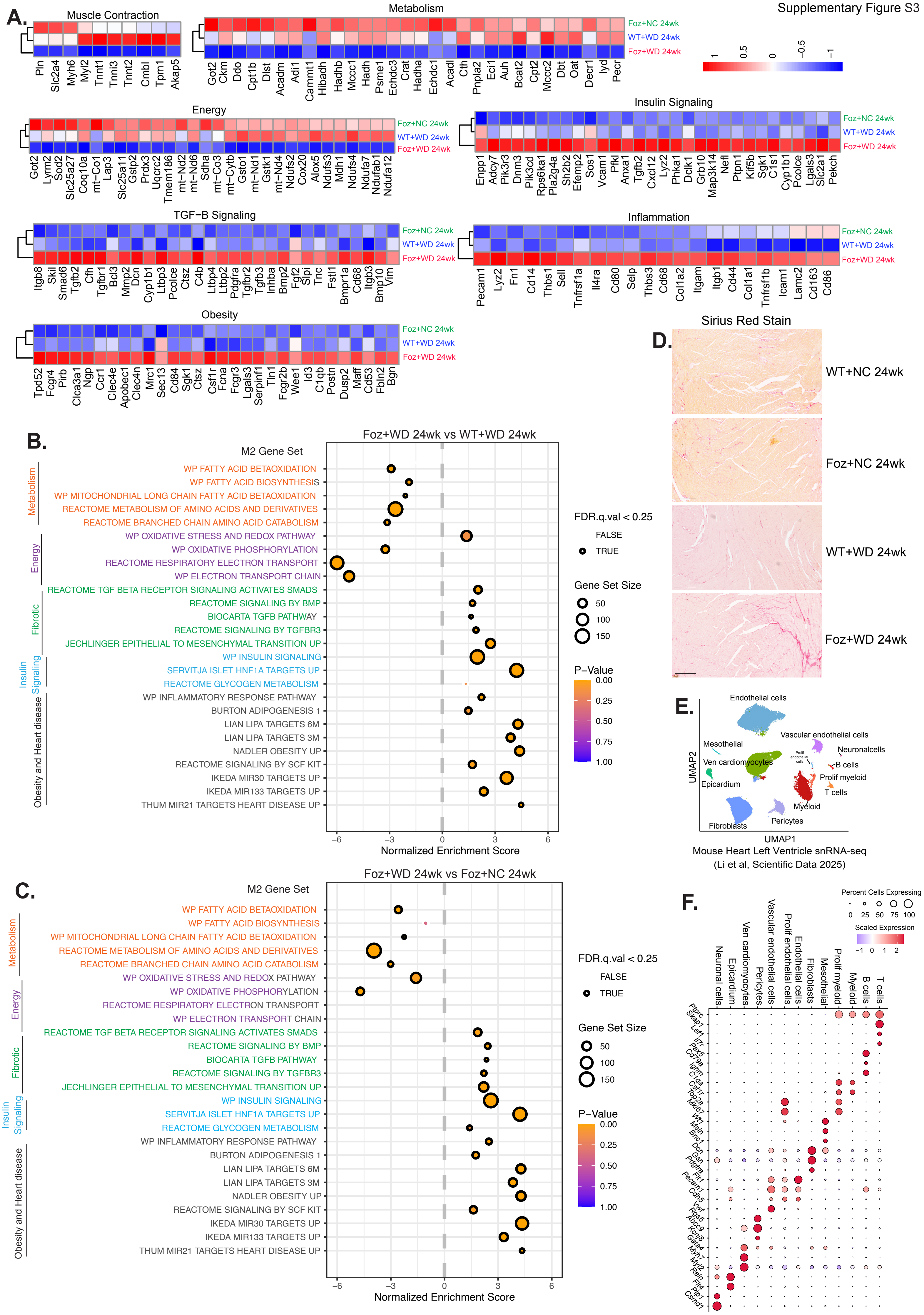

Supplementary Figure S4

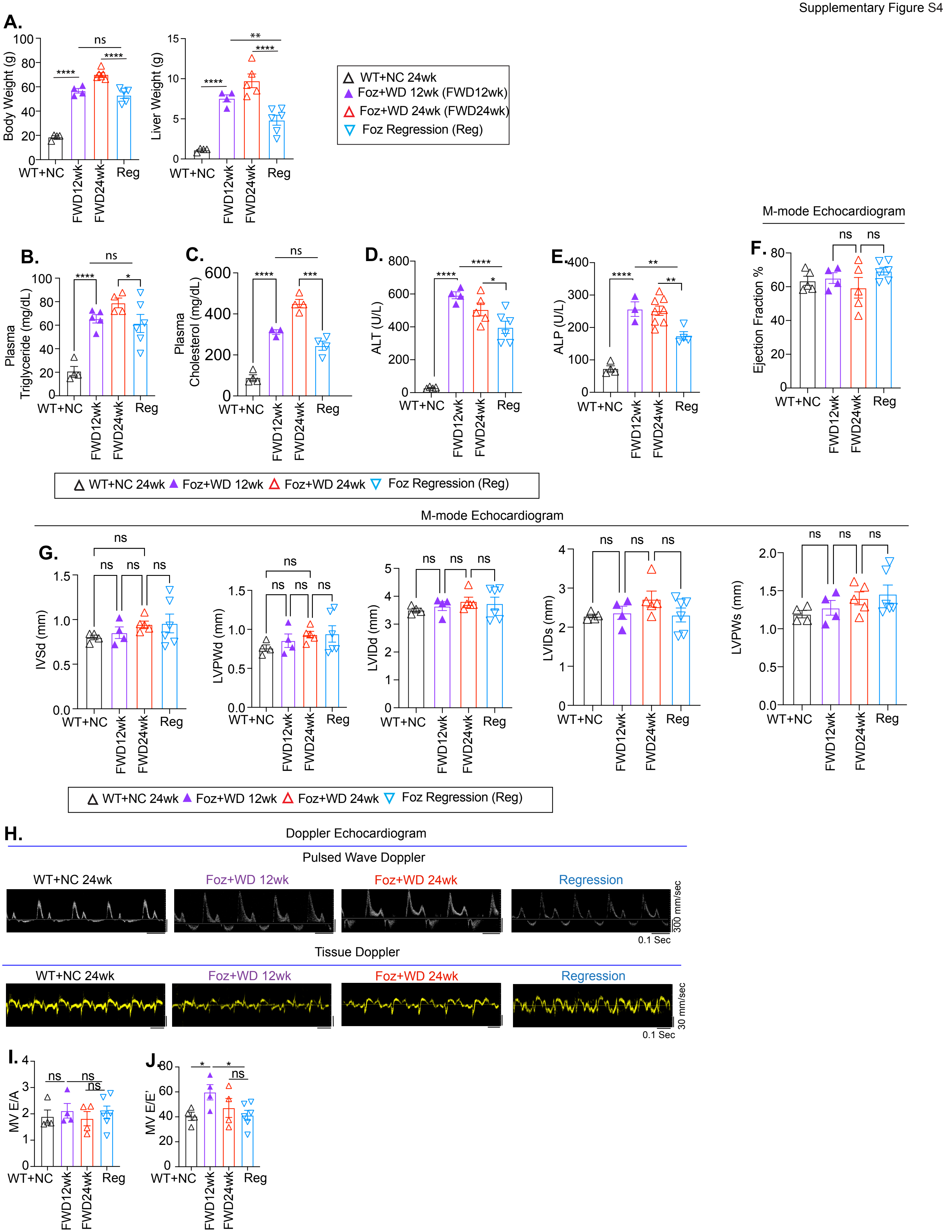

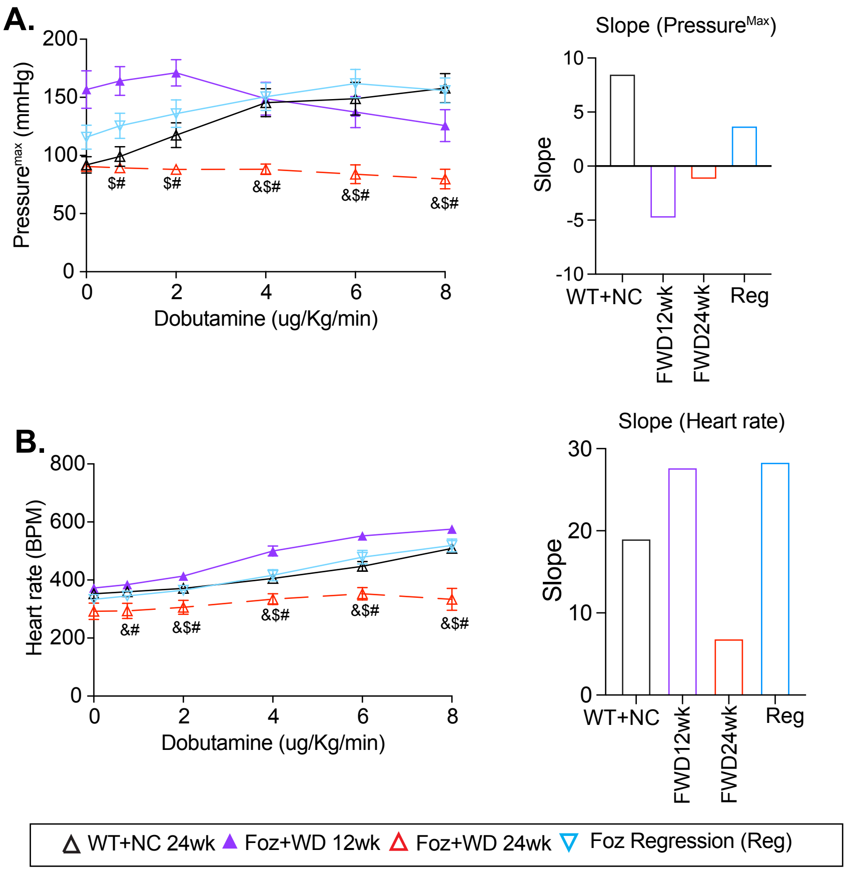


**C.**

| Group | Parameters | WT+NC 24 wk | FOZ+NC 24 wk | WT+WD 24 wk | FOZ+WD 12 wk | FOZ+WD 24 wk | REGRESSION |
|---|---|---|---|---|---|---|---|
| | Obesity | Normal | High | High | High | Very high | High |
| | Plasma cholesterol | Normal | High | High | High | Very high | Improved |
| | Plasma triglyceride | Normal | High | High | High | High | Improved |
| Metabolic | Fasting blood glucose | Normal | High | High | High | High | Improved |
| | Fibrosis (SR+ area) | Normal | Low | Low | High | Very high | Improved |
| | Fibrosis (METAVAIR staging) | Normal | F0-1 | F0-1 | F2 | F4 | F1 |
| | Alt | Normal | High | High | Very high | Very high | Improved |
| Hepatic | Steatosis | Normal | High | High | High | High | Improved |
| | Cardiac hypertrophy | Normal | Increased | Increased | Increased | Markedly increased | Remains high |
| | Cardiomyocyte dysfunction (contractility/relaxation) | Normal | Normal | Normal | Present | Present | Improved |
| | B-adrenergic responsiveness (max/min dp/dt) | Normal | Slightly reduced | Slightly reduced | Impaired | Severely impaired | Improved |
| | Relaxation constant (tau) | Normal | Normal | Normal | Normal | Elevated | Similar to healthy |
| | BNP levels | Normal | Normal | Normal | Increased | Markedly increased | Improved |
| Cardiac | Ejection fraction (EF) | Normal | Normal | Normal | Preserved (>50) | Preserved (>50) | Preserved (>50) |
| | Mortality | Normal | None | None | None | High | Rescued |

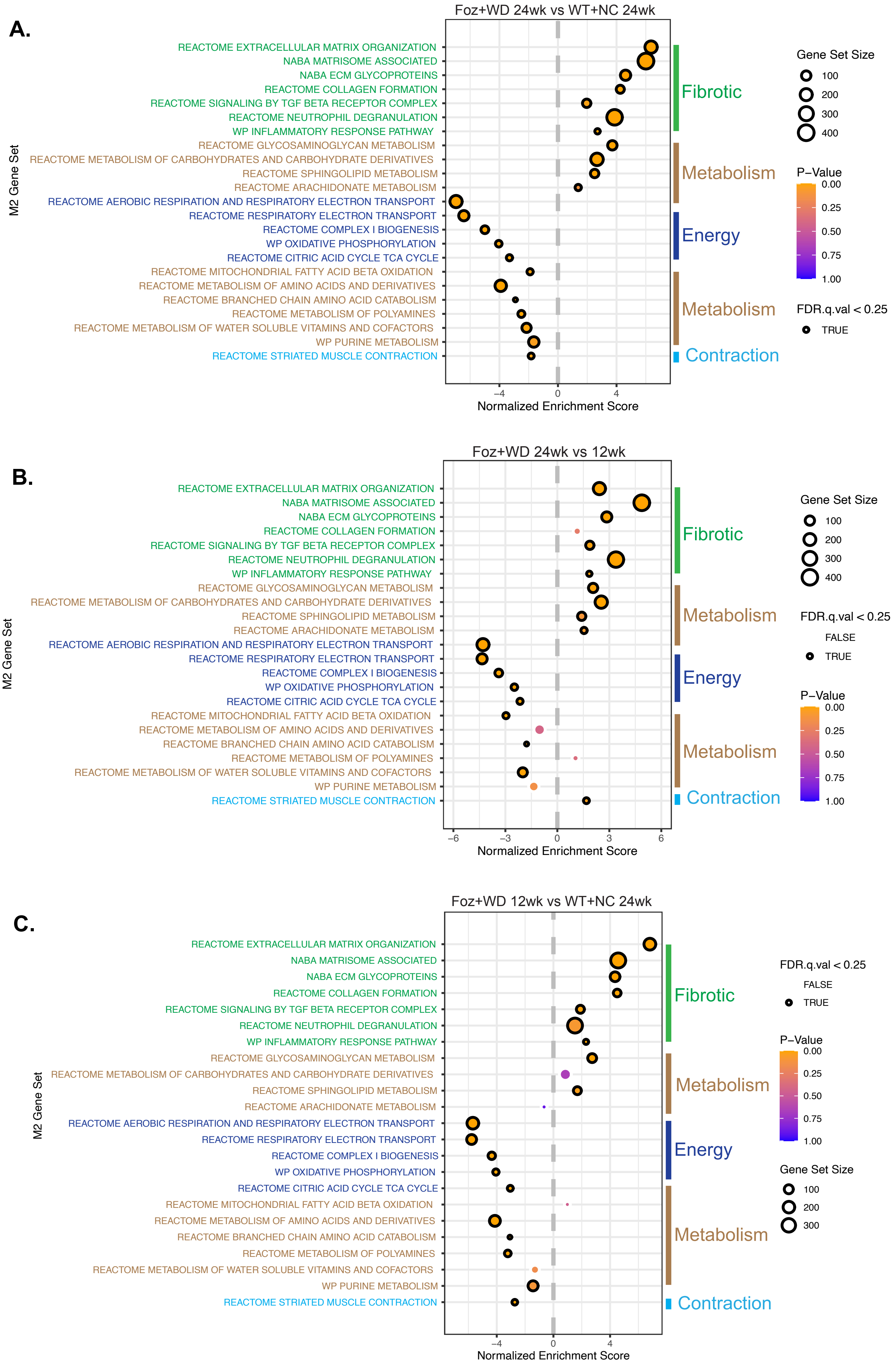
A.
Foz+WD 24wk vs WT+NC 24wk
M2 Gene Set
REACTOME EXTRACELLULAR MATRIX ORGANIZATION
NABA MATRISOME ASSOCIATED
NABA ECM GLYCOPROTEINS
REACTOME COLLAGEN FORMATION
REACTOME SIGNALING BY TGF BETA RECEPTOR COMPLEX
REACTOME NEUTROPHIL DEGRANULATION
WP INFLAMMATORY RESPONSE PATHWAY
REACTOME GLYCOSAMINOGLYCAN METABOLISM
REACTOME METABOLISM OF CARBOHYDRATES AND CARBOHYDRATE DERIVATIVES
REACTOME SPHINGOLIPID METABOLISM
REACTOME ARACHIDONATE METABOLISM
REACTOME AEROBIC RESPIRATION AND RESPIRATORY ELECTRON TRANSPORT
REACTOME RESPIRATORY ELECTRON TRANSPORT
REACTOME COMPLEX I BIOGENESIS
WP OXIDATIVE PHOSPHORYLATION
REACTOME CITRIC ACID CYCLE TCA CYCLE
REACTOME MITOCHONDRIAL FATTY ACID BETA OXIDATION
REACTOME METABOLISM OF AMINO ACIDS AND DERIVATIVES
REACTOME BRANCHED CHAIN AMINO ACID CATABOLISM
REACTOME METABOLISM OF POLYAMINES
REACTOME METABOLISM OF WATER SOLUBLE VITAMINS AND COFACTORS
WP PURINE METABOLISM
REACTOME STRIATED MUSCLE CONTRACTION
Fibrotic
Metabolism
Energy
Metabolism
Contraction
−4
0
4
Normalized Enrichment Score
Gene Set Size
100
200
300
400
P−Value
0.00
0.25
0.50
0.75
1.00
FDR.q.val < 0.25
TRUE
B.
Foz+WD 24wk vs 12wk
M2 Gene Set
REACTOME EXTRACELLULAR MATRIX ORGANIZATION
NABA MATRISOME ASSOCIATED
NABA ECM GLYCOPROTEINS
REACTOME COLLAGEN FORMATION
REACTOME SIGNALING BY TGF BETA RECEPTOR COMPLEX
REACTOME NEUTROPHIL DEGRANULATION
WP INFLAMMATORY RESPONSE PATHWAY
REACTOME GLYCOSAMINOGLYCAN METABOLISM
REACTOME METABOLISM OF CARBOHYDRATES AND CARBOHYDRATE DERIVATIVES
REACTOME SPHINGOLIPID METABOLISM
REACTOME ARACHIDONATE METABOLISM
REACTOME AEROBIC RESPIRATION AND RESPIRATORY ELECTRON TRANSPORT
REACTOME RESPIRATORY ELECTRON TRANSPORT
REACTOME COMPLEX I BIOGENESIS
WP OXIDATIVE PHOSPHORYLATION
REACTOME CITRIC ACID CYCLE TCA CYCLE
REACTOME MITOCHONDRIAL FATTY ACID BETA OXIDATION
REACTOME METABOLISM OF AMINO ACIDS AND DERIVATIVES
REACTOME BRANCHED CHAIN AMINO ACID CATABOLISM
REACTOME METABOLISM OF POLYAMINES
REACTOME METABOLISM OF WATER SOLUBLE VITAMINS AND COFACTORS
WP PURINE METABOLISM
REACTOME STRIATED MUSCLE CONTRACTION
Fibrotic
Metabolism
Energy
Metabolism
Contraction
−6
−3
0
3
6
Normalized Enrichment Score
Gene Set Size
100
200
300
400
FDR.q.val < 0.25
FALSE
TRUE
P−Value
0.00
0.25
0.50
0.75
1.00
C.
Foz+WD 12wk vs WT+NC 24wk
M2 Gene Set
REACTOME EXTRACELLULAR MATRIX ORGANIZATION
NABA MATRISOME ASSOCIATED
NABA ECM GLYCOPROTEINS
REACTOME COLLAGEN FORMATION
REACTOME SIGNALING BY TGF BETA RECEPTOR COMPLEX
REACTOME NEUTROPHIL DEGRANULATION
WP INFLAMMATORY RESPONSE PATHWAY
REACTOME GLYCOSAMINOGLYCAN METABOLISM
REACTOME METABOLISM OF CARBOHYDRATES AND CARBOHYDRATE DERIVATIVES
REACTOME SPHINGOLIPID METABOLISM
REACTOME ARACHIDONATE METABOLISM
REACTOME AEROBIC RESPIRATION AND RESPIRATORY ELECTRON TRANSPORT
REACTOME RESPIRATORY ELECTRON TRANSPORT
REACTOME COMPLEX I BIOGENESIS
WP OXIDATIVE PHOSPHORYLATION
REACTOME CITRIC ACID CYCLE TCA CYCLE
REACTOME MITOCHONDRIAL FATTY ACID BETA OXIDATION
REACTOME METABOLISM OF AMINO ACIDS AND DERIVATIVES
REACTOME BRANCHED CHAIN AMINO ACID CATABOLISM
REACTOME METABOLISM OF POLYAMINES
REACTOME METABOLISM OF WATER SOLUBLE VITAMINS AND COFACTORS
WP PURINE METABOLISM
REACTOME STRIATED MUSCLE CONTRACTION
Fibrotic
Metabolism
Energy
Metabolism
Contraction
−4
0
4
Normalized Enrichment Score
FDR.q.val < 0.25
FALSE
TRUE
P−Value
0.00
0.25
0.50
0.75
1.00
Gene Set Size
100
200
300

Supplementary Figure S7

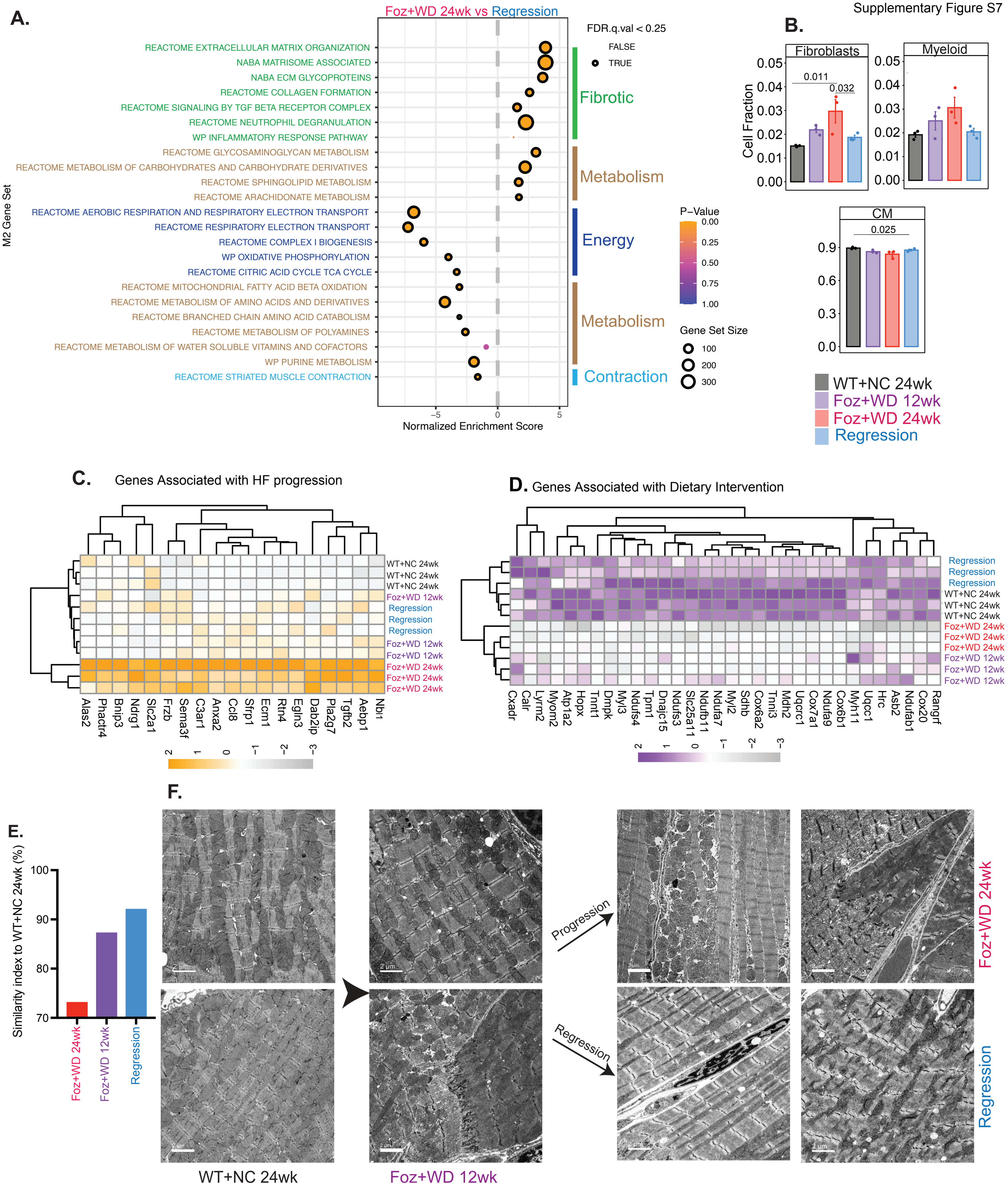
